# A New Likelihood-based Test for Natural Selection

**DOI:** 10.1101/2021.07.04.451068

**Authors:** Helmut Simon, Gavin Huttley

**Affiliations:** Research School of Biology, the Australian National University

**Keywords:** selective neutrality, site frequency spectrum, ACKR1

## Abstract

We present a new statistic for testing for neutral evolution from allele frequency data summarised as a site frequency spectrum, which we call the relative likelihood neutrality test or *ρ*. Classical methods of testing for natural selection, such as Tajima’s D and its relatives, require the null model to have constant population size over time and therefore can confound demographic change with natural selection. *ρ* can directly incorporate a null hypothesis reflecting general demographic histories. It has a natural Bayesian interpretation as an approximation to the log-probability of the null model, given the data. We use simulations to show that *ρ* has greater power than Tajima’s D to detect departure from neutrality for a range of scenarios of positive and negative selection. We also show how *ρ* can be adapted to account for sequencing error. Application to the ACKR1 (FYO) gene in humans supported previous studies inferring positive selection in sub-Saharan populations which were based on inter-population comparisons. However, we did not find the signal of selection to be maximal in the region of the FY*O or Duffy-null allele in these populations. We also applied *ρ* to investigate in greater detail a region on the 2q11.1 band of the human genome that has previously been identified as showing evidence of selection. This was done for a range of populations: for the European populations we incorporated a demographic history with a bottleneck corresponding to the putative out of Africa event. We were able to localise signals of selection to some specific regions and genes. Overall, we suggest that *ρ* will be a useful tool for identifying genomic regions that may be subject to natural selection.

## INTRODUCTION

Detecting the influence of natural selection on a genomic region provides critical insight into genomic function and the evolution of a population. For example, identifying regions where positive selection has operated can illuminate advantageous functional modifications. Genes located where negative selection against deleterious mutations occurs highlight preservation of function. The task of associating a population genetic signal with a phenotype is generally difficult, but a useful step is to refine the locus in which the signal of selection occurs. The effects of selection on patterns of variation can be difficult to distinguish from those of population demographic history (Akey et al., 2004; Sabeti et al., 2006). Approaches have been proposed to overcome this (Stajich and Hahn, 2004; Akey et al., 2004; Nielsen et al., 2009), but to date these entail some compromise of the power of the tests involved.

The site frequency spectrum (SFS) is a widely used representation of population genetic variation. For a sample of *n* orthologous sequences containing polymorphic sites, the SFS is the vector of counts, indexed by *i*, of the number of segregating sites for which mutant alleles occur in exactly *i* members of the sample. Evolutionary forces influence the distribution of variant alleles. For example, in a population whose size has steadily increased as we approach the present, a greater proportion of the current population and its ancestors will have been extant recently. A larger proportion of the mutations that occurred in the population over time will then have occurred in such more recent individuals, and will therefore have had less time to spread through the population. The result is an SFS that is skewed toward a greater number of rare variants (lower *i*) (Keinan and Clark, 2012). To take another example, variant alleles subject to weak purifying selection, and any neutral variant alleles linked to such alleles, will tend to be removed from a population before reaching higher frequencies. Consequently, purifying selection also results in a bias toward low frequency variants (Charlesworth et al., 1993; Cvijović et al., 2018). These examples demonstrate the influence that evolutionary forces can have on patterns of variation, and also that different evolutionary forces can result in apparently similar patterns of variation.

The most commonly used general-purpose test for neutrality in non-recombinant sequences remains Tajima’s D (Tajima, 1989; Vitti et al., 2013). Tajima’s D is motivated by comparing the average number of pairwise genetic differences (*π*) between individuals and the total number of segregating sites in the sample (*S*_*n*_). Under a Wright-Fisher model, the expected values of these quantities are equal, modulo a factor depending only on *n*. Tajima’s D and the related statistics of Fu and Li (1993) and Fay and Wu (2000) can be derived directly from the SFS (Achaz, 2009). Tajima (1989) calculated the variance of his statistic under a Wright-Fisher null and used a beta distribution as an approximation of its distribution. The use of the variance as a normalizing factor has generally been considered to justify the application of a consistent threshold for determination of statistical significance across a wide range of sample sizes and mutation rates.

While Tajima’s D has proved itself useful in a wide range of applications, detailed analysis has shown shortcomings. One is that Tajima’s D has low power in detecting weak selection for smaller sample sizes (Simonsen et al., 1995). Any method that aims to improve on Tajima’s D and the related statistics of Fu and Li (1993) and Fay and Wu (2000) will need to use more information from the data than is captured by the three quantities on which they depend: *S*_*n*_, *π* and the number of singletons (Simonsen et al., 1995).

Three approaches have been used to embed Tajima’s D and related statistics in methods that can distinguish the effects of selection from those of demographic change. One is based on the concept that demographic change affects all parts of a genome in the same manner, while the effect of selection is local. Regions for which, for example, the absolute value of Tajima’s D significantly exceeds the genome average are interpreted as potentially undergoing selection (for example, Hamblin et al., 2002).

This approach does not require a specific demographic history to be identified for the population, but assumes that the influences of demography and selection do not cancel each other. Another approach uses inter-population comparisons. A strong deviation from neutrality that occurs at a given locus in one population but not another may be a signal of selection at the locus in the first population (Sabeti et al., 2006). Alternatively, it potentially reflects different demographic histories. Intuitively, such a signal would be considered more powerful if it were consistent across related populations (for example a set of African populations compared to a set of European populations). Both of these approaches can complement methods for identifying selection at specific loci, as will be seen below.

A third approach assumes that a null demographic history has been identified for the population. It aims to detect loci for which the probability that some set of summary statistics derived from the data can be produced by this demographic history alone is small compared to the genome average. For example, Stajich and Hahn (2004) analyzed 151 loci in samples from both African-American and European-American populations. They used a consensus demographic scenario, incorporating a bottleneck corresponding to migration from Africa, as a null hypothesis for the European-American population. They performed a large number of simulations of this scenario to estimate the expected value of a set of summary statistics (in this case *π, θ* and Tajima’s D) under this hypothesis. They then calculated this set of summary statistics for the 151 loci and identified those loci for which these statistics were most different to the results of the simulations. A constant population size Wright-Fisher null was used in a similar way for the African-American population. A similar approach is taken in Akey et al. (2004) using a different set of summary statistics, including Tajima’s D and those of Fu and Li (1993) and Fay and Wu (2000). In this case, demographic parameters for a null model were selected by a grid search matching summary statistics from coalescent simulations using these parameters with statistics calculated from the empirical population. Another related approach was taken by Nielsen et al. (2009). They considered a group of 13400 protein coding genes, taken from a sample of 20 European-American and 19 African-American genomes. They then identified those loci that were outliers in the group in terms of various metrics based on their SFS, such as bias toward low, high or intermediate frequency variants, or in terms of the probability of the SFS under a multinomial model where the probability parameters are the average allele frequencies across the group (G2D test). All three approaches aim to identify regions that are outliers relative to the overall genome in some relevant way. While an outlier approach may identify a subset of candidate loci potentially enriched for selection (Kelley et al., 2006), it does not of itself provide a sufficient basis to assess the probability of a null model being the one responsible for the pattern of variation at a given locus.

While the G2D test of Nielsen et al. (2009) uses a type of likelihood function, it is a quasi-likelihood under a multinomial model whose probability parameter is an average of the empirical allele frequencies. This “multinomial-mean” likelihood is a zero-order approximation to the true likelihood, which depends on the full distribution of allele frequencies defined by genealogical trees generated by an evolutionary model.

Our approach is based on two general principles. The first is that using likelihood is the best way to maximize the information obtained from the SFS data. We therefore define an algorithm to approximate the likelihood of the null model. Secondly, to assess the probability that a null model is responsible for the data requires a comparison to the likelihood of alternative hypotheses, as is implied by Bayes’ theorem. We develop a method to approximate the probability distribution function of models given the data, and evaluate it at the null model. We show that this is equivalent to calculating the ratio of the likelihood of the null model to the expected likelihood over the space of all evolutionary models. We represent the latter space by zero-order approximations similar to those used in the method of Nielsen et al. (2009) referred to above. We refer to our approach as the relative likelihood neutrality test and define a test statistic *ρ*. We show using simulations that it typically has greater power in detecting selection, and apply the method to scanning several regions of the human genome for evidence of selection.

## MATERIALS AND METHODS

In this section, we will define a test for neutrality using observed data in the form of a SFS. The SFS for a sample of *n* aligned sequences was defined previously as the vector **s** of numbers *s*_*i*_ for *i* = 1, …, *n* −1, where *s*_*i*_ is the number of segregating sites for which the mutant allele occurs in *i* members of the sample. Since we assume the infinite-sites model of mutation (Kimura, 1969), no more than one mutant allele can occur at any site. Calculation of the SFS also requires knowledge of which segregating allele is the mutant, that is, we require an outgroup. We will derive a way to express the likelihood of observing a given SFS in terms of parameter definitions that relate mutation and genealogy.

### Computing likelihoods of models

To estimate the probability of a particular SFS, we make use of the “backwards in time” perspective on population genetics reflected in coalescent theory (Kingman, 1982a,b). This is illustrated in Figure 1(a) for a present day sample of *n* = 6 non-recombinant sequences. In principle, we can derive the genealogy of a sample by tracing back the ancestry of each member of the sample until the first time that two members share a common ancestor (members a and b in Figure 1(a)). At this time we say that a “coalescent event” has occurred and the number of ancestors of the original sample is reduced to *n* − 1. Iterating this step for *n* − 1 coalescent events, we arrive at the single most recent ancestor of the entire sample (at the top of Figure 1(a)). As shown in Figure 1(a), the genealogy then takes the shape of a tree. (We assume that all samples in all generations experience a coalescent event after some finite number of generations, but that only one pair of ancestors of the sample experience a coalescent event in any single generation.) We can characterize the tree by its combinatorial structure and a sequence of *n* − 1 branch lengths which separate the coalescence times. We denote these lengths by *t*_*k*_, where for 2 ≤*k* ≤ *n, t*_*k*_ is the length of time (number of generations) for which there are *k* ancestors of the sample. The combinatorial structure refers only to the ancestry relationship between sequences and their ancestors.

**Figure 1.**
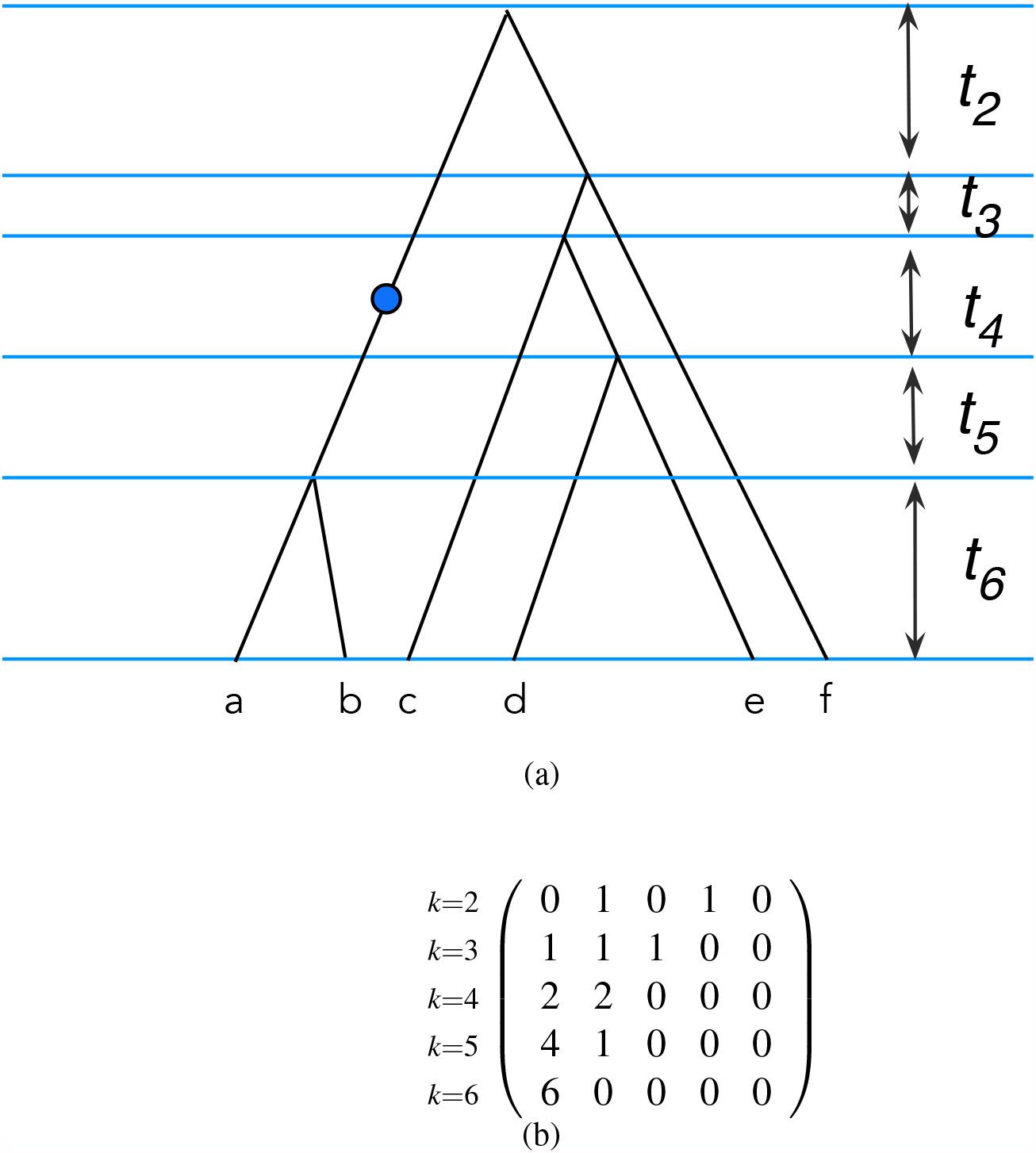
Representation of genealogical relationships. (a) The genealogy of a sample of six sequences. Time flows from the top of the figure to the bottom. The bottom is the time at which the samples were taken. The intervals between the times of coalescence events (marked by horizontal blue lines) are usually referred to as branch lengths and denoted by *t*_*k*_ for 2 ≤ *k* ≤*n*. The index *k* denotes the number of branches in the tree during each interval.The blue circle represents a mutation event occurring in the time period during which the sample has four ancestors. (b) The matrix representation of the combinatorial structure of the tree. Matrix rows correspond to levels of the tree as denoted by *k*. Matrix columns correspond to the number of descendants of a branch present at each value of *k*.

The SFS from a sample is determined by the genealogical tree associated with the sample and the mutation events that occurred within that genealogy. That is, the number of members of the sample in which the mutation is found can be identified by tracing the descendants of the mutated sequence through the tree. For example, in Figure 1(a) the mutation (shown as a blue circle) occurred when *k* = 4 and has two present-day descendants, a and b, in the sample. For each *k*, we can define a partition of the *n* sequences in the present-day sample into *k* blocks by assigning all descendants of each branch present at the time during which there are *k* ancestors of the sample to a single block. This yields an unordered *k*-partition of *n* for each *k* with 2 ≤*k* ≤*n*. For example, when *k* = 4 for the sample depicted in Figure 1(a), the 4-partition of the sample is {{a,b}, {c}, {d,e}, {f}} from which we derive the integer sequence {2, 1, 2, 1}, where each value is the size of the partition block. In this way, we can associate each tree with a sequence of unordered *k*-partitions of *n* for *k* = 2, …, *n*. We refer to this sequence as the *tree partition class* (TPC) of the tree. We further transform each unordered 4-partition into an ordered vector of length 4, where the positions of the vector correspond to the block sizes 1, 2, 3, 4. Our example above can thus be represented as the vector (2, 2, 0, 0, 0), indicating there are 2 blocks of size 1, 2 blocks of size 2 and 0 of sizes 3, 4 and 5. Successive such vectors for *k* = 2, …, *n* can be configured as a matrix which can then be identified with the TPC of a genealogy and provides a useful representation of the genealogy (Figure 1(b)). The rows correspond to *k*, while the columns correspond to the size of partitions. We denote the matrix derived in this way from a tree *ψ* by *A*_*ψ*_. For a tree with *n* leaves, *A*_*ψ*_ is an (*n* − 1) × (*n* − 1) matrix. The matrix entry *A*_*ψ*_ [*i, j*] gives the number of branches at level *k* = *i* + 1 of the tree *ψ* with *j* descendants. This corresponds, in our earlier terminology, to the number of partitions of the *n* leaves with block size *j* induced by the *k* = *i* + 1 nodes at the *k*^*th*^ level.

The following fundamental identities follow immediately for any row *i* (1 ≤*i* ≤ *n* −1) in *A*_*ψ*_ :

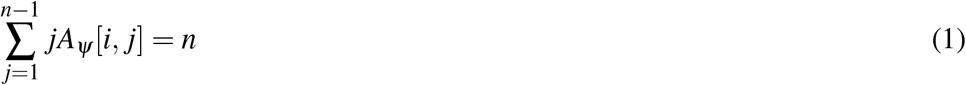

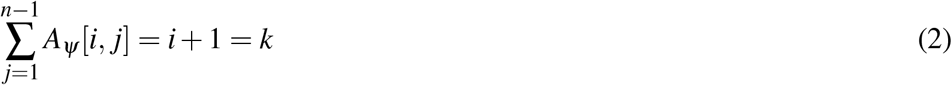

We denote the TPC of the tree *ψ* by 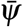 and, more generally, the set of TPCs for a given *n* by 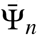. Each TPC corresponds to a way of recursively transforming a set of *n* unlabelled objects (the root node of a tree) into *n* singletons (the external nodes of the tree) via a sequence of *n* − 1 binary partitions. At each step, a partition of the *n* objects into *k* blocks is transformed into a partition into *k* + 1 blocks by dividing one of the *k* blocks (necessarily one with more than one element) into two. The cardinalities 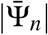 are finite and comprise the integer sequence A002846 in Sloane et al. (2003). The set 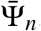 is equivalent to the set ℱ_*n*_ used to represent genealogical trees in Sainudiin et al. (2011) and the matrices *A*_*ψ*_ correspond to Sainudiin’s *f* -matrices. As the matrix *A*_*ψ*_ depends only on the TPC, then 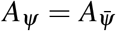. The TPC equivalence relation is not the same as topological equivalence. It is clear that trees that are topologically equivalent can belong to different TPCs, as topology does not take account of the temporal sequence in which bifurcations occur. The converse is also true, and an example is given in Supplementary Figure S1.

We can define a probability measure on 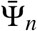. A standard assumption of coalescent models is that all pairs of individuals present at a point in time are equally likely to be the next to coalesce. Viewed forward in time, this is equivalent to requiring that all the possible binary partitions at each step are equally probable. If we label the original *n* objects and place them in a defined order, there are *n* − 1 gaps between them. Then, the recursive subdivision of the partitions described above is equivalent to the selection of a non-repeating sequence of these *n* − 1 gaps, which can in turn be represented by a permutation of the integers 1, …, *n* − 1. The outcome is a sequence of ordered partitions, which can be transformed into a set of unordered partitions by “forgetting” the order. If we count the instances of TPCs for all possible permutations of 1, …, *n* − 1, we obtain a measure on 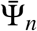 that reflects our assumption that all pairs of sequences in a sample are equally likely to be the next to coalesce. This measure is analogous to that resulting from a Yule process on trees and we refer to it as the Equal Rates Markov (ERM) measure on 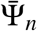 (Mooers and Heard, 1997). *We denote the probability of a TPC* 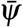 under this measure by 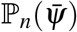.

The SFS summarises the numbers of descendants of mutations segregating in a sample. For the genealogy *ψ*, the probability that a mutation occurring while the sample has *k* ancestors has *j* descendants is given by *A*_*ψ*_ [*k* −1, *j*]*/k*. The expected value of this probability integrated over all TPCs using the ERM measure is denoted by *p*_*n,k*_(*j*) and is given by the following formula (Fu, 1995; Griffiths and Tavaré, 1998):

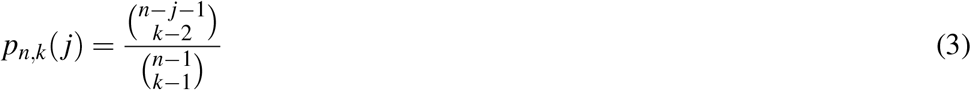

If *j > n−k* + 1, *p*_*n,k*_(*j*) = 0, consistent with the observation that a node at level *k* can have no more than *n* − *k* + 1 descendants. Note that we have from the so-called “hockey stick” identity on Pascal’s Triangle (DeTemple and Webb, 2014, Theorem 1.38):

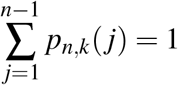

We define the expected value *A*_*n*_ of the matrices 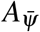 under the ERM measure on 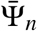 by:

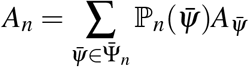

For a given *n*, the value of *A*_*n*_ can be calculated from equation (3):

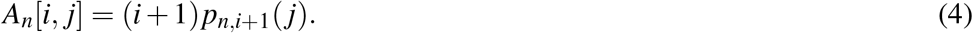

*A*_*n*_ is a triangular matrix and it can be seen from equation (4) that the diagonal entries are positive. Hence it is invertible (Bhaskar and Song, 2014). A formula for the inverse is given in Supplementary Information Section 2.

We derive a likelihood function for the SFS, conditioned on a tree *ψ* with branch lengths *t*_*k*_ for 2 ≤*k* ≤*n*. The total length of branches at level *k* having *j* descendants is 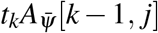. Then, summing over *k* gives the total length of branches in *ψ* in which any mutation will have *j* descendants, i.e.:

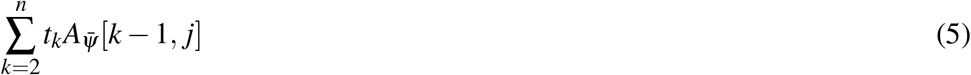

The sum of all these components is necessarily equal to the total tree length:

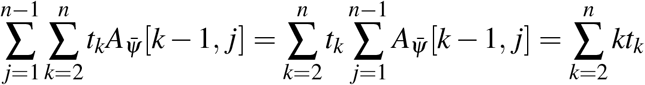

where the last equality follows from equation (2).

Our assumption of the infinite sites theory of mutation (Kimura, 1969) requires that the timescales we are considering are sufficiently small relative to the mutation rate that each mutation occurs at a unique segregating site. We assume that the number of mutations occurring in a single branch is given by a Poisson distribution with parameter given by the product of branch length (in generations) and the mutation rate. Thus, if we denote by *q* _*j*_ the probability that a mutation in the tree has *j* descendants in the sample, we can calculate *q* _*j*_ as the proportion of the total tree length composed of branches with *j* descendants (equation 5), i.e.:

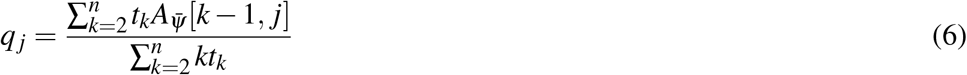

We can represent this in vector notation as:

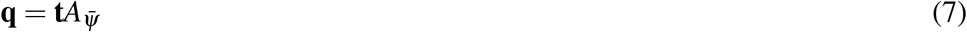

where 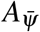 is the matrix representation of the TPC of the tree *ψ*; **t** denotes the (*n* − 1) row vector of branch lengths relative to total tree length, i.e.

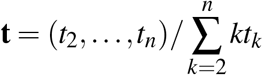

and **q** is the vector of probabilities *q*_1_, …, *q*_*n*−1_ from equation (6).

Our observed data is the SFS, namely **s** = (*s*_1_, …, *s*_*n*−1_). It follows from the definition of the SFS that:

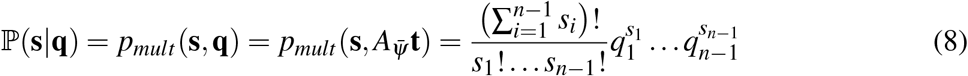

where *p*_*mult*_ is the probability mass function of the multinomial distribution, conditioned on *n* −1 (the number of event categories) and *S*_*n*_ (the number of draws). Equation (8) defines the likelihood function for our model. Each vector **q** defines a multinomial probability distribution on the SFS. Since the vectors **q** are determined by genealogical trees, they are generated by an evolutionary model that generates trees, (e.g. Wright-Fisher).

We now define a representation of the Wright-Fisher model and calculate its likelihood for a given SFS. This model, **M**_0_, is assumed to have specific values of the sample size *n* and the number of segregating sites *S*_*n*_. Equation (8) shows how to calculate the probability of the SFS **s** for a given tree matrix 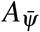 and vector of relative branch lengths **t**. The probability distribution on 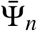 under any model **M** that we consider will be the ERM measure. We denote the probability distribution of relative branch lengths under the model **M** by *T*_*n*_. We further assume the combinatorial structure of trees, represented by the tree matrix, is independent of the branch lengths. Then we can use the Law of Total Probability to calculate the likelihood of *s* under the model **M** by taking the expectation over the joint (Cartesian product) distribution 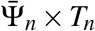:

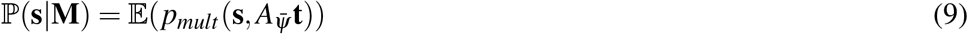

For the Wright-Fisher model, it is known (Hein et al., 2005) that if **T**_*n*_ = (*T*_2_, …, *T*_*n*_) represents the joint distribution of branch lengths, then **T**_*n*_ is a sequence of independent exponential distributions with mean 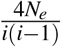 where *N*_*e*_ is the diploid effective population size, i.e.:

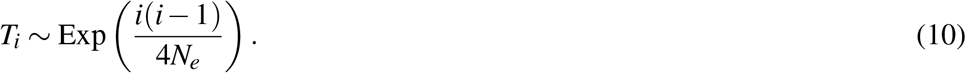

We can evaluate the right hand side of equation (9) by Monte Carlo integration. That is, we take *G* samples from 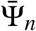 using the ERM measure and from **T**_*n*_ using equation (10) and multiply them to get *G* variates *q*_1_, …, *q*_*G*_ (equation 7). Using equation (8), we then estimate ℙ (**s**|**M**) by:

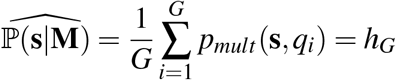

Standard results for Monte Carlo integration (Robert et al., 2010, p. 65) also allow us to estimate the standard error *sde*_*G*_ by:

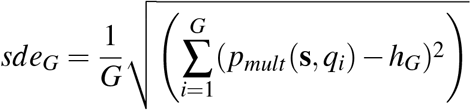

### Calculating the likelihood of more complex demographic models

We can extend this method to calculate the likelihood of a model corresponding to a demographic history other than constant population size. In order to define a model reflecting variable population size, we use the approach referred to earlier of modifying the Wright-Fisher model to accommodate a variable population size by transforming the time variable to reflect the local rate of change in population size (Slatkin and Hudson, 1991; Griffiths and Tavaré, 1994, 1998). This method is the one used in popular coalescent simulation software packages such as Hudson’s `ms` (Hudson, 2002). We could in principle use such simulation software directly to generate samples of branch lengths which are used in the method described above. Using a full coalescent simulation package for this purpose is generally too inefficient to be practical, particularly for larger sample sizes. We addressed this by restricting ourselves to a simple piece-wise constant population model. In such a model, we specify a set of time points *τ*_0_, …, *τ*_*m*_ going backward in time from the present (*τ*_0_), together with a population size value for each interval from *τ*_*i*_ to *τ*_*i*+1_. We then generate Wright-Fisher branch variates using equation (10) as before and transform them to generate samples of the distribution of branch lengths corresponding to our piece-wise constant population model. This functionality is available in the selectiontest library described below.

### An evolutionary model can be characterised by a probability distribution

An evolutionary model **M** can be pictured as a method of generating random TPCs (or tree matrices) and branch length vectors or, equivalently, defining probability distributions on these objects. It follows from equation (7) that the model **M** also results in a probability distribution, which we denote by **Q**, on the probability vectors **q**. To generate SFS instances from the probability distribution **Q**, we first sample vectors **q** from **Q** and then sample from the multinomial distribution with parameters **q** and *S*_*n*_ (the number of draws). This is analogous to using a population genetic simulation package which first stochastically generates trees and then stochastically adds mutations to each tree. The likelihood of the model depends only on **Q** as, by integrating equation (8) over **Q** we have:

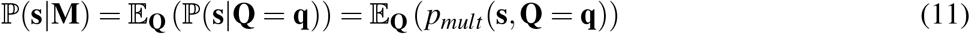

### Estimating the probability of a model, given observed data

We wish to estimate the probability that **s** was produced by a specific model **M**_0_ of an evolutionary process as this represents the most direct measure of the explanatory power of this model for the data. (For example, **M**_0_ may be the Wright-Fisher model, with a specific sample size *n* and number of segregating sites *S*_*n*_.) To be precise, we will estimate the value of the conditional probability density function (PDF) *f* (**M**_0_ |**s**) where *f* is the prior PDF on the space of evolutionary models. We will not assume any prior knowledge of the model that produced the data and reflect this by assuming that the prior distribution *f* is uniform, with *f* (**M**) being constant. Using Bayes’ theorem we have:

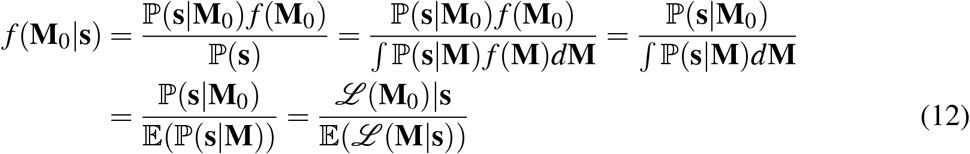

In the third equality we have used the fact that *f* (**M**) = *f* (**M**_0_) for all **M** to factor out these terms.

We see that *f* (**M**_0_ | **s**) is equivalent to the ratio of the likelihood of the null model to the expected value of the likelihood taken over all possible models. We now estimate *f* (**M**_0_ |**s**) using equation (12). We have shown earlier in this section how to estimate the numerator, that is, the likelihood ℒ (**M**_0_ | **s**) = ℙ (**s** |**M**_0_). We next consider the estimation of the denominator 𝔼(*ℒ* (**M** | **s**)) = 𝔼(ℙ (**s** | **M**)). As we do not have a definition of the space of all evolutionary models, we take the heuristic approach of representing each model **M** by the mean probability vector 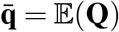, where **Q** is the probability distribution corresponding to **M**. We next determine the space of probability vectors that correspond to valid evolutionary models in this way. If we denote the components of 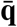 by 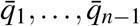, the value of 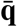 for an arbitrary evolutionary model is given by (Griffiths and Tavaré, 1998, equation 3.3):

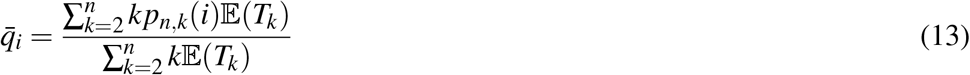

In this equation, an evolutionary model results in probability distributions *T*_*k*_ on branch lengths with expected values 𝔼(*T*_*k*_). The TPCs have been “integrated out” using the ERM measure, hence the presence of the coefficients *p*_*n,k*_(*i*) from equation (3). The left hand side 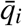 represents the mean probability that a mutation has *i* descendants averaged over the distributions of branch lengths and tree structures. Using equation (4), we can rewrite equation (13) as:

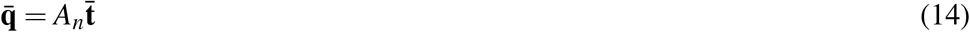

The valid values for the (mean) relative branch lengths 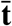 are the elements of the (*n* − 2)-dimensional simplex in 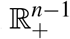 given by:

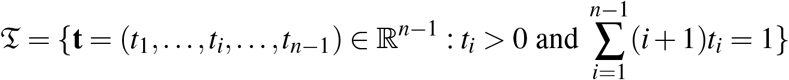

From equation (14), vectors **q** that correspond to valid relative branch lengths and hence to evolutionary models are points in the image *A*_*n*_(𝔗). *A*_*n*_(𝔗) is a simplex which is a subset of the standard simplex Δ^*n*−2^ in ℝ^*n*−1^. We will denote the uniform distribution on *A*_*n*_(𝔗) by 𝒰.

Having represented each model by a probability vector 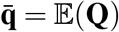, we approximate the distribution **Q** by the multinomial distribution with parameters 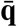 (vector of probabilities) and *S*_*n*_ (number of draws). We then substitute these approximations into equation (11) to obtain:

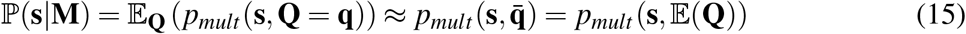

The approximation of the likelihood in this way is similar to the use of a multinomial distribution on averaged empirical allele frequencies in the composite likelihood approach used by Nielsen et al. (2009).

To estimate 𝔼(ℙ(**s** | **M**)), the denominator in equation (12), we take the expected value of both sides of the approximation in equation (15). For both sides, we integrate over the uniform distribution, giving:

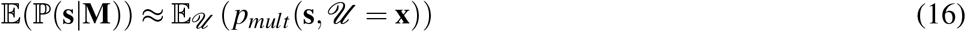

We evaluate the right hand side of equation (16) by Monte Carlo integration. To do this requires a method to sample from 𝒰. We use the observation that 𝔗 is the image of Δ^*n*−2^ under **J**_*n*_, where **J**_*n*_ is the diagonal matrix whose diagonal elements are 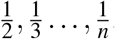 Since *A*_*n*_ and **J**_*n*_ are both bijections, *A*_*n*_ **J**_*n*_ is a linear bijection from Δ^*n*−2^ to *A*_*n*_(𝔗). That is, all maps in the following diagram are bijections.

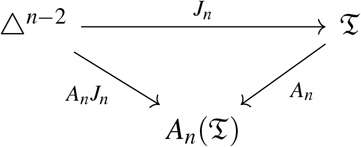

Since the linear transformation of a uniform distribution is uniform (Rudin, 1987, Theorem 2.20), we can sample uniformly from *A*_*n*_(𝔗) by sampling points *x* uniformly from Δ^*n*−2^ using a Dirichlet distribution whose parameters are all equal to one, and transforming them into points *A*_*n*_*J*_*n*_*x*. We thus have a Monte Carlo algorithm for estimating 𝔼(ℙ(**s**|**M**)) using equation (16).

### Testing for neutrality

We use the value of the conditional PDF at the null model, as calculated using equation (12), as a measure of the support of the data for the null model **M**_0_. We showed above how to estimate the numerator of the right hand side of equation (12), the likelihood of the model given the data, using equation (9). We also showed how to estimate the denominator, using equation (16). We thus define the following test statistic for the null hypothesis corresponding to the model **M**_0_:

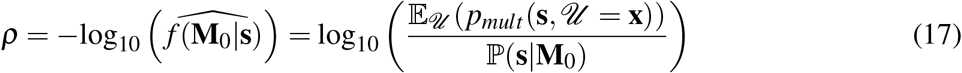

Thus *ρ* can be interpreted in terms of value of the conditional PDF at the null model and also as the log-ratio of the expected likelihood over all models to the likelihood of the null model. *ρ* has been designed to accord with the conventions for statistics based on likelihood ratios: that a logarithm is taken and that the likelihood of the alternate model is the numerator. Thus, larger values of *ρ* represent a decreasing support for the null model.

The statistic *ρ* can be adapted for use as a hypothesis test by comparing its value to that for data produced by the neutral model using the same parameters, *n* and *S*_*n*_. For hypothesis testing, thresholds need to be defined to achieve a given level of significance or *p*-value, usually denoted by *α*. The level of significance corresponds to the false positive rate (FPR) or Type I error rate (the rate of rejection of cases that are in fact neutral) that we are prepared to accept. In this paper, we will use a significance value of *α* = 5%, consistent with the investigations of Tajima’s D and similar tests of neutrality by Simonsen et al. (1995) and Braverman et al. (1995). Thresholds are calculated by performing multiple simulations of a Wright-Fisher model for these parameters, calculating *ρ* for each SFS data set produced and identifying the desired quantile, 5% in our case. Thresholds need to be calculated for specific values of *n* and *S*_*n*_. Table 1 shows estimated threshold values for *ρ* for an FPR of 5%, for a range of values of *n* and *S*_*n*_. If *n* and *S*_*n*_ are sufficiently large, the probabilities involved are frequently too low to allow a quantile to be calculated. For this reason, no result is shown for the case *n* = 100 and *S*_*n*_ = 100.

**Table 1.**
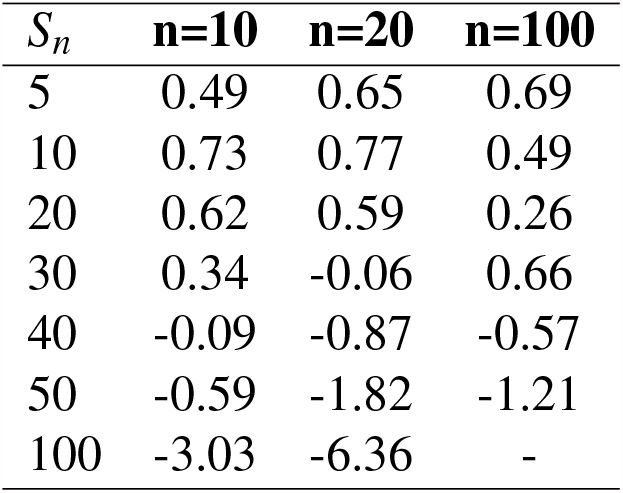
Thresholds used for *ρ* to test the hypothesis of neutrality at an FPR of 5% for the values of *n* and *S*_*n*_ shown. The number of simulations run in each calculation is ≥ 200000.

### Data

Variant Call Format (VCF) data for the human genome was downloaded from the 1000 Genomes Project (1KGP) Phase 3 (1000 Genomes Project Consortium; Auton et al., 2015), which uses GRCh37 coordinates. The populations are listed in Supplementary Information Section 3. For the analysis of the human chromosome band 2q11.1 region, the NCBI Build 34 (UCSC hg16) coordinates used by the HapMap project (chr2:96250000-96750000) were converted to GRCh37 coordinates using pyliftover, a Python implementation of UCSC LiftOver (Tretyakov, 2013). We only used one chromosome per sampled individual to minimize the effect of diploidy on sampling error. Details of populations analysed are given in the Supplementary Information.

Chimpanzee VCF data used in investigation of the ACKR1 gene (in Discussion) was downloaded from the Great Ape Genome Project (Prado-Martinez et al., 2013). There was missing data in the form of samples in which allele information was not provided for some variants. This was addressed by excluding all such variants. Macaque VCF data for the the ACKR1 gene was sourced from the Macaque Genotype and Phenotype Resource (Bimber et al., 2019). Samples with missing data were excluded from the analysis.

### Software

Software modules to allow users to apply the methods described in this paper were written in Python version ≥ 3.7 and are available in the Python library selectiontest. The functionality provided includes calculation of *ρ*; calculation of thresholds for *ρ*; sampling from **Q** for the Wright-Fisher model, the uniform model and models involving piece-wise constant demographic histories; sampling from the space of tree matrices 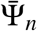 according to the ERM probability measure; calculation of Tajima’s D from the SFS; and the calculation of an SFS from variant data in VCF format. Functionality is made available both as Python modules and as a command line interface. See documentation at https://selectiontest.readthedocs.io/ for details.

For low values of *n* and *S*_*n*_ it is possible to validate the calculation of the likelihood ℒ (**M**_0_ |**s**) = ℙ(**s**| **M**_0_) by comparing the results to the long run frequencies of SFS configurations generated by repeated simulations. This was done for the case where **M**_0_ is the Wright-Fisher model; *n* = 6 and *S*_*n*_ = 4, which yields a manageable 70 SFS configurations (results not shown).

Scripts and Jupyter notebooks developed to perform analyses specific to this work, such as comparisons of *ρ* and Tajima’s D, were also written in Python version ≥ 3.7 and are freely available under the GPL at https://github.com/helmutsimon/NeutralityTest and https://zenodo.org/record/4679991.

The simulated data sets used for this paper were produced by the software packages msms 3.2rc-b163 (Ewing and Hermisson, 2010) and fwdpy11 0.9.0 (Kelleher et al., 2018). msms implements an extension of the Hudson coalescent simulation `ms` software (Hudson, 2002) and was used to simulate positive selection. msms is a Java program which we called from within a Python script. fwdpy11 is a Python library for forward simulation of population genetic models. It was used to simulate negative and background selection. Forward simulation is intrinsically less efficient than coalescent-based (backward) simulation.

Other software dependencies include `cogent3` 2019.12.6a (Knight et al., 2007), pyliftover 0.3 (Tretyakov, 2013), scipy 1.2.1 (Virtanen et al., 2020), msprime 0.7.0 (Kelleher et al., 2016), numpy 1.16.3 (Virtanen et al., 2020), pandas 0.24.2 (McKinney, 2010), pysam 0.15.0, (Li et al., 2009), pyvcf 0.6.8, (Casbon, J. et al., 2012), click 6.7 (Ronacher, 2009), scitrack 0.1.3 (Huttley, 2016), matplotlib 3.0.3 (Hunter, 2007) and seaborn 0.9.0 (Waskom et al., 2017).

### Data availability statement

The authors state that all data necessary for confirming the conclusions presented in the article are represented fully within the article. Supplementary Information, preprocessed data and scripts and Jupyter notebooks developed specifically to perform the data sampling and analyses reported in this work are available at Zenodo https://zenodo.org/record/4679991, doi 10.5281/zenodo.4679991 under the Creative Commons Attribution-Share Alike license.

## RESULTS

### Hypothesis testing using simulated data

We tested the performance of *ρ* in detecting natural selection using simulation studies for two scenarios: positive selection and purifying (including background) selection. We used Tajima’s D (hereafter simply referred to as “D”) as a benchmark due to its widespread use and its preeminence among related statistics (Simonsen et al., 1995). As D tests deviation from a Wright-Fisher population, we adopt the null hypothesis of a Wright-Fisher model to make comparisons objective. Two sets of simulations were carried out. We first conducted multiple simulations of the null Wight-Fisher model to generate an empirical distribution of SFS data. This data was used to determine the thresholds for *ρ* and D corresponding to the desired significance level. For *ρ* the calculated thresholds are shown in Table 1 and for D at Supplementary Information Section 5. The second set of simulations synthesised SFS data using the alternate model. From these we determined the power of each statistic, that is, the proportion of cases for which the null hypothesis is correctly rejected at the calculated thresholds.

Semi-dominant positive selection was simulated over a range of selection coefficients, *s*, using the msms software (Ewing and Hermisson, 2010). As mutant alleles that have even a significant selective advantage behave similarly to neutral alleles when they are present at low frequencies (Kimura, 1983, §8.4) we do not expect to observe any signature of selection in such a case. Accordingly, tests for positive selection are most effective at loci where the selected allele has reached or is near to fixation in the population. We conditioned the simulation on the allele under selection reaching fixation 4000 generations before the present. We also conditioned the simulations on 10 segregating sites (*S*_*n*_ = 10). Alignments with a relatively low number of segregating sites are of interest as they allow the investigator to test for selection in narrower genomic segments.

A difference between *ρ* and D is that D has lower and upper thresholds (is two-sided) while *ρ* is one-sided. Values of D are negative if there are a greater number of low-frequency or high-frequency variants than would be expected under neutrality (Braverman et al., 1995), and are positive in the opposite case. In the case of positive selection, one expects a greater relative frequency of rare alleles due to the removal of older variants of intermediate frequency by selective sweeps (Kelley et al., 2006), resulting in a negative value for D. If one is seeking evidence only for positive selection based on a skew toward low frequency alleles, it is arguable that values exceeding the upper threshold (of ∼ 2) do not count as successes. We thus consider D as a one-sided statistic and attach the significance level (5% in our case) only to the lower tail of the distribution. (The authors are not aware of D being used in this way elsewhere in the literature.) Results for two-sided thresholds are also included because one may not be able to anticipate the nature of the selective process, if any, that has given rise to the data. An example of this latter case is given by the scan of a region of human chromosome band 2q11.1 discussed below. These considerations do not apply to *ρ*, which has a single threshold, representing a degree of divergence from the null.

We found that *ρ* showed greater power than D across the range of selection coefficients (*s*) and sample sizes (*n*) for both one-and two-sided tests (Figure 2). Even considering D as a one-sided statistic with a significance level of 5%, *ρ* still outperforms it for lower levels of selection (*s* ≤ 10^−5^) and to some extent for low sample sizes (Figure 2(a)). These results confirm previous observations that D is relatively weak in detecting selection when sample sizes are low (Simonsen et al., 1995). However, for the larger sample size of *n* = 100, the performance of the two statistics is quite close. An exception occurs for the case *n* = 10, for which D gives a substantially better outcome than *ρ* for the relatively large selection coefficient of 5 × 10^−3^, while the power of *ρ* is less for *s* = 5 × 10^−3^ than for *s* = 10^−3^.

**Figure 2.**
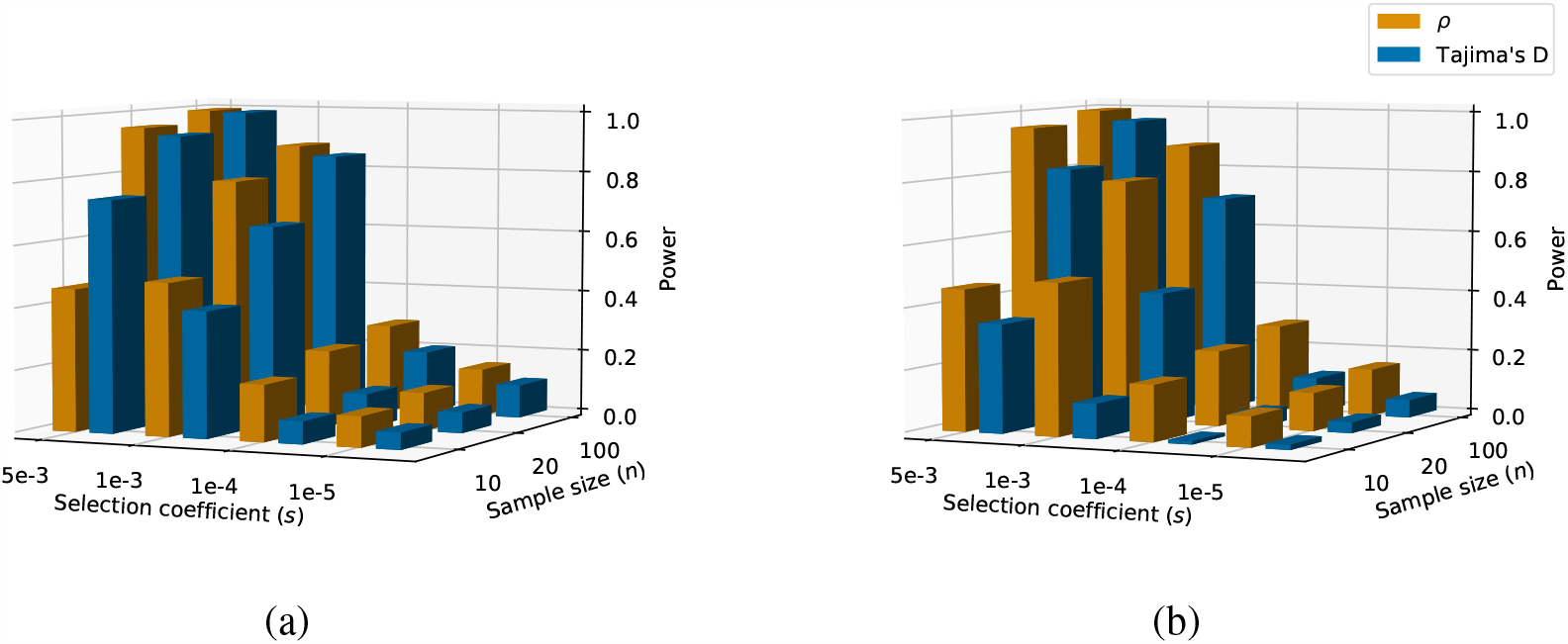
Testing for neutrality using *ρ* and Tajima’s D for data simulated by models of positive selection. The height of the bars indicates the power of the respective tests (the proportion of times that the a test correctly rejected the hypothesis of neutrality at a significance levels of 5%). (a) A one-sided test for D. (b) A two-sided test for D (see text). Simulations were conditioned on *S*_*n*_ = 10. The number of SFS configurations generated for each model was 10000.

We also simulated negative selection, involving mutations that are mildly deleterious and hence have a selection coefficient *s <* 0. Unlike the case of positive selection, mutant alleles do not reach fixation under a model of negative selection. Instead, the models invoke recurrent mutation to the negatively selected allele. Provided that |*sN*_*e*_ |*≫* 1, where *N*_*e*_ is the effective population size, these mutant alleles are generally eliminated from the population. In addition, neutral variants that are linked to the mildly deleterious alleles will also tend to be eliminated from the population. This effect on neutral alleles is referred to as background selection (Charlesworth et al., 1993), typically involving selection coefficients −.05 *< s <* 0. Thus negative selection also has characteristic effects on allele frequencies as observed in the SFS of sampled loci. As mentioned above, mildly deleterious alleles behave similarly to neutral alleles at low frequencies, so the proportion of both selected and linked neutral variants at these low allele frequencies are expected to be similar to those of the neutral model. At intermediate allele frequencies, both deleterious and linked neutral variants will be affected. A large proportion of deleterious variants will be directly eliminated from the population, depressing the proportion of variants at intermediate frequencies. A proportion of neutral variants linked to deleterious mutations will also be eliminated. If recombination is absent or negligible, the effective population size for neutral variants is reduced by a factor of *e*^*u/*2*hs*^ where *u* is the per genome per generation mutation rate of deleterious mutations and *h* is the dominance coefficient (Charlesworth et al., 1993). Thus background selection similarly lowers the proportion of neutral variants at intermediate allele frequencies. The strength of background selection on neutral variants is determined by the (positive) parameter *λ* defined by:

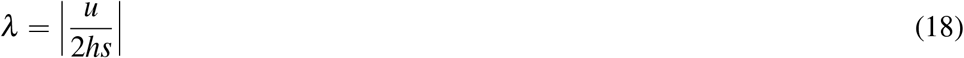

The larger *u* is relative to *hs*, the greater the proportion of sequences carrying deleterious mutations that will survive long enough to accumulate neutral mutations, increasing the strength of background selection. A more detailed analysis of the effect of background selection on allele frequencies of linked neutral variants is provided in Cvijović et al. (2018). For our purposes, it is important to note that in the vicinity of a locus at which deleterious mutations occur, there are effects on both the deleterious mutations themselves and on linked neutral variants, driven by two parameters: *s* and *λ*. The relative effect of these will to some extent be influenced by the boundaries of the genomic region chosen for analysis.

We simulated loci subject to recurrent neutral and deleterious mutations in the absence of recombination using the forward simulation package *fwdpy11* (Kelleher et al., 2018). As was the case for positive selection, we could consider D as a one sided statistic if we are looking for evidence that the subject population has a greater proportion of low frequency alleles than would be expected under neutrality. Results for both one-sided and two-sided tests using D are shown in Figure 3, varying the parameters *s* and *λ*, while holding *n* = 20. We included both neutral and deleterious mutations as, when testing actual genomic regions, we do not generally have *a priori* knowledge of where or indeed whether deleterious variants occur. We see that *ρ* is again systematically more effective than D for the case of negative selection, even though the average number of segregating sites in each simulation is large, varying between 65 and ∼ 400.

**Figure 3.**
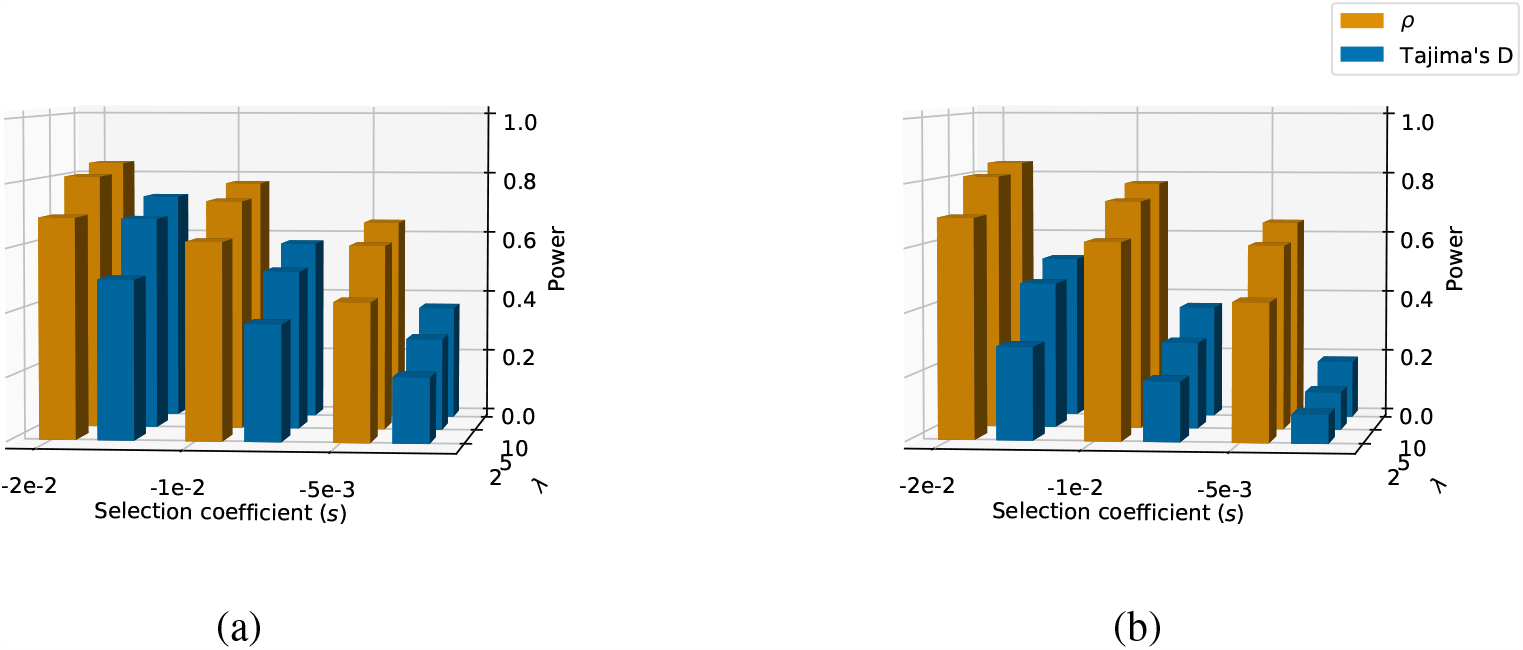
Testing for neutrality using *ρ* and Tajima’s D for data simulated under models of negative selection. The parameters are selection coefficient (*s*) and the ratio of selected mutation rate to selection coefficient (*λ* -see equation (18)). The height of the bars indicates the power of the respective tests (the proportion of times that the test correctly rejected the hypothesis of neutrality at a significance levels of 5%). (a) A one-sided test for D. (b) A two-sided test for D (see text). In all cases the sample size *n* = 20, the diploid population size was 1000 and the neutral mutation rate is equal to the selected mutation rate. The thresholds for *ρ* and D are conditioned on *n* and *S*_*n*_. The number of SFS configurations generated for each model was 10000.

One can also use *ρ* to detect demographic change in the absence of selection. In Figure 4 we show results from simulations of exponential growth for a range of growth rates and sample sizes (*n*), comparing *ρ* to D. As expanding populations also result in an excess of rare variants, we again show one-sided and two-sided thresholds for D. In these examples, we include a relatively large sample size *n* = 250. Again, *ρ* generally shows greater power to detect deviation from the Wright-Fisher model. The exception occurs for the largest sample size *n* = 250. Here we see that for all population growth rates, the power of *ρ* is less than for *n* = 100. This is most likely due to numerical problems arising from underflow errors due to probabilities of specific SFS data sets being extremely low for large *n*. Further simulations have shown that some more complex demographic scenarios, such as population bottlenecks, can confound *ρ*, causing it to have power less than 50% (data not shown). In such cases, the null hypothesis is not rejected, even though the evolutionary model is very different from the null.

**Figure 4.**
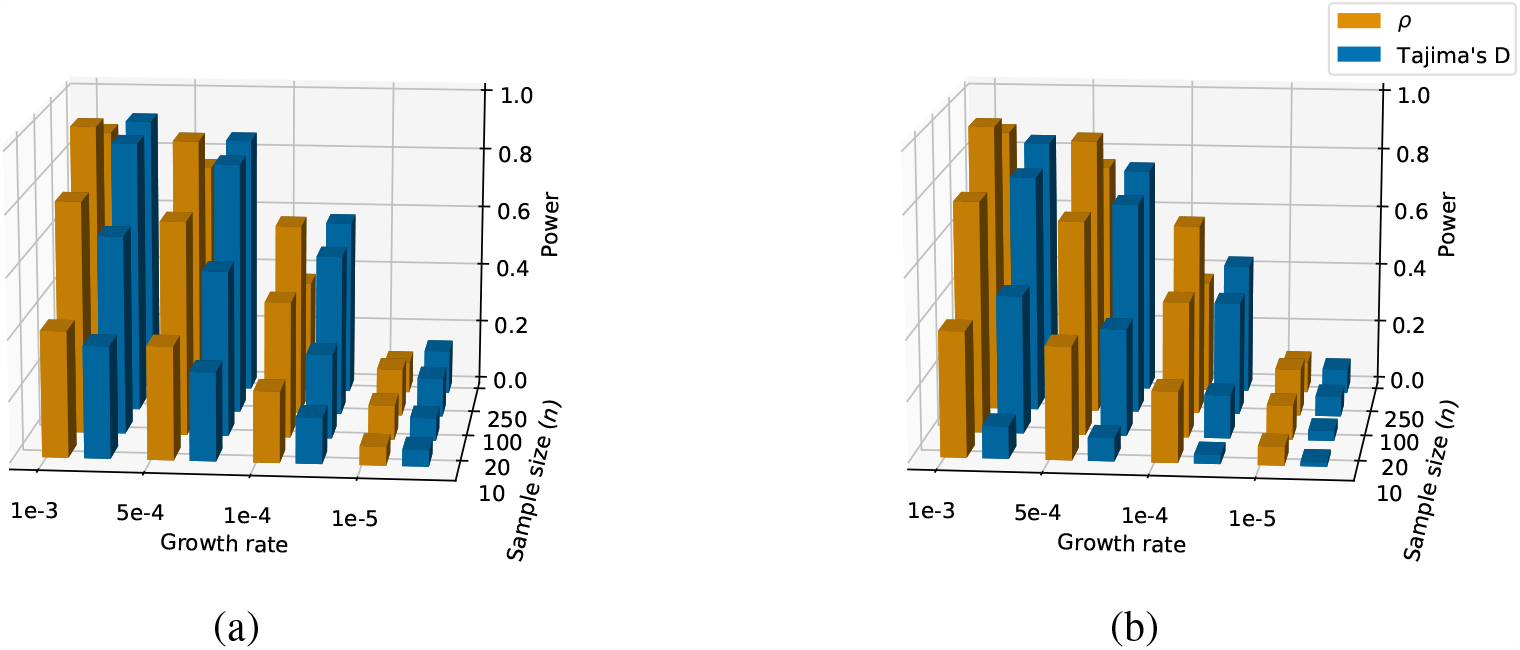
Testing for neutrality using *ρ* and Tajima’s D for data simulated by models of exponential population growth. The parameters are per-generation growth rate and sample size (*n*). The height of the bars indicates the power of the respective tests (the proportion of times that the a test correctly rejected the hypothesis of neutrality at a significance levels of 5%). (a) A one-sided test for D. (b) A two-sided test for D (see text). In all cases the number of segregating sites *S*_*n*_ = 10. The thresholds for *ρ* and D are conditioned on *n* and *S*_*n*_. The number of SFS configurations generated for each model was 10000.

### Signals of selection in *ACKR1* (the Duffy antigen gene)

The *ACKR1* gene encodes the Duffy antigen, which is expressed on the surface of both red and white blood cells and is necessary for the invasion of red blood cells by the vivax malaria parasite, *Plasmodium vivax* (Miller et al., 1976). *P. vivax* is one of four malarial parasites found in humans, the greatest mortality being associated with *P. falciparum* (Livingstone, 1984). *The ACKR1* gene exhibits one of the highest levels of genetic variance between human populations of any locus. The variant rs2814778, which characterizes the FY*O or Duffy-null allele, is near fixation in sub-Saharan populations, but rare elsewhere (Hamblin et al., 2002; Hodgson et al., 2014). The FY*O allele deactivates expression of the Duffy antigen and thus confers resistance to vivax malaria. This has been taken as evidence of positive selection for this allele in sub-Saharan Africa (Hamblin and Di Rienzo, 2000; Hamblin et al., 2002; Carter, 2003). This hypothesis has been questioned due to the generally inverse geographic relationship between the prevalence of vivax malaria and the FY*O allele, evidence for the origin of vivax malaria in east Asia and the low mortality associated with vivax malaria (Livingstone, 1984; Escalante et al., 2005). Genome-wide surveys using population genetic tools such as Tajima’s D have not identified this gene as a locus of selection (Hamblin and Di Rienzo, 2000; Sabeti et al., 2006).

We used *ρ* to investigate whether this inability to detect a departure from neutrality was caused by a lack of power in Tajima’s D and other tests. Initially, we examined all populations in the 1KGP data set (see Materials and Methods and Supplementary Information Section 3) to determine which showed a signal of selection using *ρ* over the gene region (chr1:159173097-159176290, GRCh37 coordinates). This led to samples with *n* ∼ 100 and *S*_*n*_ ∼10, so we used a 5% FPR threshold of *ρ >* 0.5 (see Table 1). We identified a signal of selection in three African populations, three east Asian populations and a population from Barbados of African descent, which for convenience we will also refer to as African. Of these seven populations, the FY*O variant was found to occur in all the African population samples and none of the Asian samples.

A specific model for selection at the Duffy-null allele in African populations was proposed in Hamblin et al. (2002). The model likelihood approach can be adapted to compare this specific hypothesis to a neutral hypothesis. The scenario in Hamblin et al. involved an effective population size *N*_*e*_ ∼ 8000, a (homozygote) selection coefficient of 0.002 and fixation occurring ∼ 33000 years ago. To calculate likelihoods of this model, we need samples of the associated allele frequency probability vectors **q**, using the sample sizes *n* corresponding to the data from the four African populations which show evidence of selection over the gene region: GWD; LWK; ACB and ESN. We simulated the model using msms. As msms does not output branch lengths, we approximated a sample from the distribution of probability vectors by setting a large value of *S*_*n*_ and normalizing the SFS values generated, which yields maximum likelihood estimates for the multinomial parameters **q**. We then compared the likelihoods of the Hamblin model for the SFS instances for these four populations and the region chr1:159173897-159176290 to the likelihoods of a neutral model. This resulted in odds ratios ranging from 5.3 (LWK) to 6.8 (ACB) favoring selection according to the Hamblin model over a neutral model. This may be regarded as “substantial” evidence for selection at the *ACKR1* gene in African populations, using the scale proposed in Jeffreys (1998, Appendix B).

In order to see whether the signal of selection could be localized to the neighbourhood of the rs2814778 variant, we divided the gene locus into four segments of ∼800 bp. To ensure sufficient segregating sites in each segment, we pooled all the African and all the east Asian populations (not only those showing positive selection.) The results are shown in Table 2. For these samples, *n* ∼ 600 and *S*_*n*_ ∼ 5 −10, resulting in a 5% threshold of *ρ >* 0.1. On this basis, *ρ* gives a signal of non-neutrality in all regions for both the African and east Asian populations, but the signal is weakest in the region chr1:159173897-159174697, in which the rs2814778 variant is located. For the pooled Asian population sample, which does not contain the variant, we see the strongest signal of non-neutrality in this region. Tajima’s D does not exceed the commonly used threshold | D | *>* 2 in any of these cases. However, simulations show that 5% two-sided confidence limits on Tajima’s D for *n* ∼ 100 and *S*_*n*_ ∼ 10 are (− 1.64, 1.97), so on this basis Tajima’s D indicates an excess of low frequency variants in most cases.

**Table 2.** Values of *ρ* and Tajima’s D calculated over 800 bp segments of the *ACKR1* gene for pooled African and east Asian 1KGP populations. The threshold for *ρ* corresponding to rejection of the null hypothesis at a significance value of 0.05 is *ρ >* 0.1. The corresponding limits for Tajima’s D are (−1.64, 1.97). Values rejecting neutrality given these thresholds are marked with an asterisk.

| Segment start | Africa |  | Asia |  |
| --- | --- | --- | --- | --- |
| | $\rho$ | Tajima's D | $\rho$ | Tajima's D |
| 159173097 | 2.54* | -1.89* | 1.08* | -1.69* |
| 159173897 | 0.52* | -1.74* | 3.08* | -1.89* |
| 159174697 | 2.65* | -1.78* | 1.03* | -1.36 |
| 159175497 | 2.21* | -1.61 | 1.24* | -1.55 |

### Analysing the relative strength of selection over a genomic region

A region in the 2q11.1 band of the human genome was identified as a candidate locus for a selective sweep in an African-American population by the HapMap project (Altshuler and Donnelly, 2005, Supplementary Table 4). The region exhibited low heterozygosity and a bias to alleles of low frequency. At that time about ∼ 6 genes were annotated in the region, without any of these being specifically identified as causing the signal of selection (Sabeti et al., 2006). The number of annotated genes in the region has now increased to ∼ 16. We used *ρ* to localise the signal of selection within the overall region and thus to filter the list of genes or other features that could be responsible. We downloaded data from the 1KGP data set (see Materials and Methods and Supplementary Information Section 3) for 5 African populations, 4 European populations, 2 American populations of African ancestry and an American population of European ancestry. For the European populations and the American population of European descent we applied a null demography involving a bottleneck corresponding to the putative migration of European populations from Africa, taken from Stajich and Hahn (2004) (details in Supplementary Information Section 4). For the African populations and the two populations of African descent from the Americas, we used a constant population size model (Stajich and Hahn, 2004). For convenience, we refer to these two groups as the European and African population groups.

We used *ρ* directly to compare the evidence for selection across different genome segments and populations, on the basis of its derivation (subject to approximation) as the value at the null model of the probability density function of evolutionary models given the data. We divided the region into 25 segments of length 20 kb and calculated *ρ* for each population in each segment. This led to sample sizes of ∼100 and ∼20 − 60 segregating sites. Based on Table 1, we would regard *ρ >* 0.7 as evidence for rejecting neutrality at a 5% significance level.

Strong signals of selection were evident across the region for the majority of populations. Consistent with the HapMap report (Altshuler and Donnelly, 2005), these signals appear strongest in the African populations. In interpreting Figure 5 it is necessary to bear in mind the effect of linkage. That is, both positive and negative selection may impact nucleotide diversity at nearby sites and this effect may extend over some distance, depending on the local recombination rate (Charlesworth et al., 1993; Innan and Stephan, 2003; Josephs and Wright, 2016). A potential illustration of this effect is particularly noticeable for population GWD in the sub-region around 96805244-97085244. This linkage effect is a significant impediment to narrowing the genomic region in which selection is occurring. As this effect will be eroded by recombination, it should be more evident in its impact on instances of more recent selection, where there has been less time for recombination to have an effect. We next examine some segments in detail.

**Figure 5.**
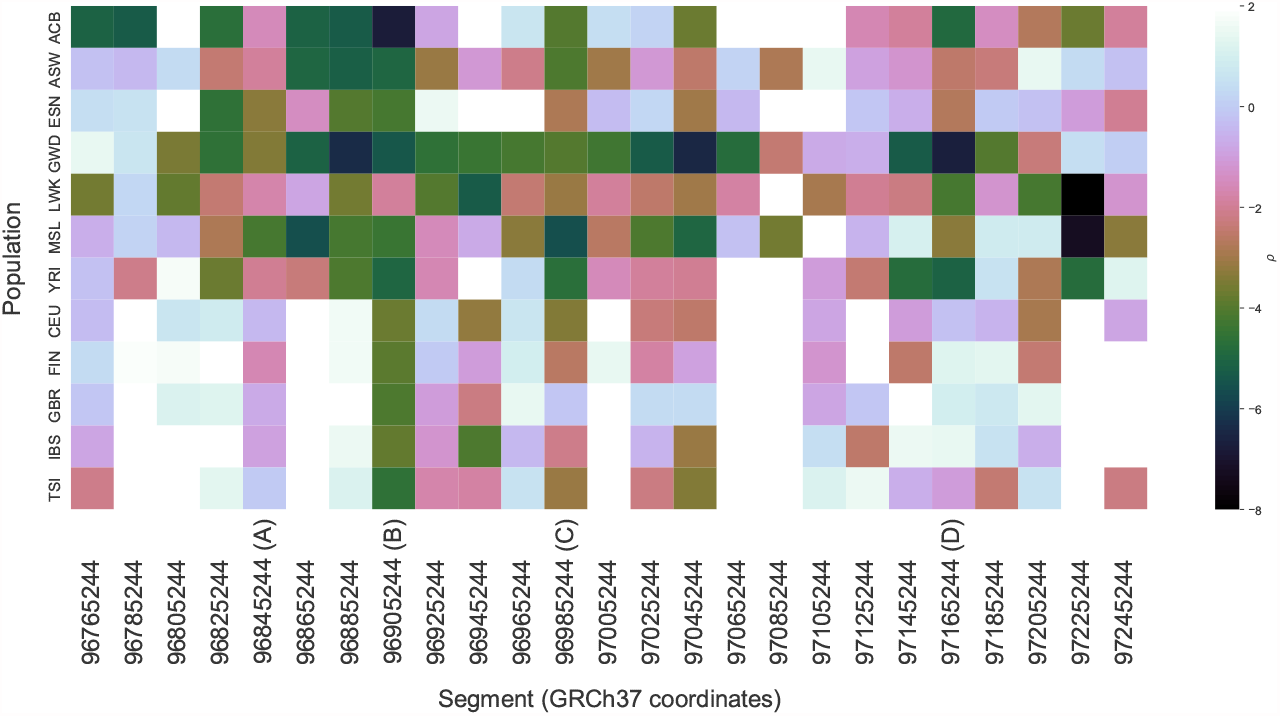
Values of *ρ* by 20-kb segment and population for the 2q11.1 region.

### chr2:96845244-96865244 (A)

This segment shows a signal of selection in both European and African populations. In the European populations *ρ* is significantly greater than in adjacent segments, while this is not apparent in African populations. The signal in European populations relative to neighbouring segments is not apparent when Tajima’s D is used (see Supplementary Figure S2). This segment overlaps with the protein-coding gene *STARD7*. An expanded ATTTC insert located within Intron 1 of this gene is associated with Familial Adult Myoclonic Epilepsy (FAME) (Corbett et al., 2019). For a sample African population (GWD), the value of *ρ* at this intron is 4.2, well in excess of the relevant 5% significance threshold of 0.7. For a sample European population (CEU), the value of *ρ* at this intron is 0.6, which is approximately equal to the relevant 5% significance threshold of 0.5. The signal of selection in this intron, which is stronger in African populations than in European populations, may result from negative selection against the ATTTC insertion.

### chr2:96905244-96925244 (B)

This segment presents a strong signal of selection in both European and African populations. It overlaps two genes, *STARD7-AS1* and *TMEM127. STARD7-AS1* is a noncoding gene with a HERV-W insert which is related to proteins involved in lipid transfer and proteolytic activity (Grandi et al., 2016). It also extends over the upstream 20 kb region. This locus shows a strong signal of selection in the GWD population (*ρ* = 10.2) but not in the CEU population ((*ρ* = − 3.7). This is consistent with Figure 5 in that a signal of selection is shown in the upstream segment for African population only.

#### TMEM127

is a transmembrane-encoding gene identified as a suppressor of pheochromocytomas, rare tumours of the adrenal system which can present before age 40 years and are associated with potentially lethal cardiovascular dysfunction (Lenders et al., 2005). For a sample African population (GWD), the value of *ρ* at the TMEM127 locus is 3.9, well in excess of the relevant 5% significance threshold of -0.6. For a sample European population (CEU), the value of *ρ* at this gene locus is 2.9, also in excess of the relevant 5% significance threshold of 0.3. Nonsense mutations in *TMEM127* have been associated with pheochromocytomas (Qin et al., 2010) and the strong signal of selection at this locus may result from background selection against such deleterious mutations.

### chr2:96985244-97005244 (C)

This segment shows a clear signal of selection in both African and European population groups. The segment overlaps with genes *AC021188*.*1, ITPRIPL1* and *NCAPH*. In an attempt to identify more precisely the locus or loci associated with selection, we performed a further analysis dividing this segment into 2-kb sub-segments. In order to obtain a sufficient number of segregating sites in each sub-segment, we pooled the African populations. This led to a sample size of 661 and around 10 segregating sites per segment. The calculated 5% FPR threshold for these parameters is approximately *ρ >* 0.1. The 2-kb sub-segments with the highest value of *ρ* started at locations 96987244 and 97003244 with values of *ρ* at 2.8 and 2.9 respectively (results not shown). The sub-segment starting at 97003244 overlaps the gene *NCAPH*, but does not overlap any exons. As it is adjacent to the boundary of the region we are considering and the signal may be caused linkage to variants outside that region, we do not consider it further. The segment starting at 96987244 overlaps with the gene *AC021188*.*1* and in particular the non-coding exon with Ensembl stable ID ENSE00001666599. It also contains numerous regulatory region variants associated with the promoter with Ensembl stable ID ENSR00000120257 and gene targets ENSG00000230747, SNRNP200, CIAO1, and ITPRIPL1.

### chr2:97165244-97185244 (D)

This segment shows a clear signal of non-neutrality in the African populations, but not in most European populations. This segment overlaps with the *NEURL3* gene only (among genes annotated in Ensembl). For a sample African population (GWD), the value of *ρ* at the NEURL3 locus is 2.0, in excess of the relevant 5% significance threshold of 0.7. For a sample European population (CEU), the value of *ρ* at this locus is also 2.0, also in excess of the relevant 5% significance threshold of 0.5. The *NEURL3* gene encodes the neuralized E3 ubiquitin protein ligase 3 (NEURL3), which inhibits hepatitis C virus (HCV) infection. Knockout of this gene has been shown to promote HCV infectivity (Zhao et al., 2018). Similarly to the case with the *TMEM127* gene, it is plausible that the signal of non-neutrality detected at this locus is also caused by negative selection operating against deleterious mutations to this gene.

## DISCUSSION

We have introduced a new statistic for testing for selective neutrality from the SFS, which we have called the relative likelihood neutrality test or *ρ. ρ* was derived from an approximation to the log posterior probability density function of evolutionary models given the observed data, evaluated at the chosen null model. *ρ* can also be interpreted as the log-ratio of the average likelihood of all evolutionary models to the likelihood of the null model (equation 17). *ρ* makes the fullest possible use of information contained in the observed SFS by its derivation from model likelihood, which captures all information in the data relevant to the explanatory power of one model relative to another. Importantly, *ρ* can test directly against null models based on any desired demographic history. This allows one to control for demographic history when testing for natural selection: the null hypothesis will be rejected when the probability that the full site frequency spectrum can be produced by the posited demographic history alone is low. This is a stronger condition than is used in the methods of Akey et al. (2004) and Stajich and Hahn (2004), which reject the null hypothesis when the probability that some set of summary statics derived from the SFS can be produced by the demographic history is low. Consequently, our method should result in a lower incidence of Type II errors relative to the null hypothesis. We demonstrated the use of a specific hypothesised null demographic history in our analysis of the 2q11.1 chromosomal region.

We used simulations to evaluate the statistical power of *ρ* in scenarios involving both positive and natural selection. For the necessarily finite number of scenarios considered, further simplifying assumptions were used, for example that recombination is absent or negligible in the locus under consideration and that all deleterious mutations in the scenario of negative selection had the same mutation rate. For these selected scenarios we carried out hypothesis testing at a 5% significance level. We used Tajima’s D as a benchmark and set out to to make the comparison as fair as possible, for example by calculating precise significance thresholds for D and by using Tajima’s D as a one sided test where we were looking solely for evidence of an excess of low frequency variants. Under most scenarios considered, *ρ* outperformed Tajima’s D, particularly in cases where selection was difficult to detect because of low sample size or low selection coefficients. Numerical issues impact the calculation of *ρ* when sample size and/or the number of segregating sites are very large (as a rule of thumb, *>* 100 in either case). In such cases, the probabilities of most specific SFS configurations are so small that their calculation is subject to underflow errors.

To test the hypothesis of neutrality for a given population using *ρ*, we need to calculate an appropriate threshold value to achieve the desired significance level (false positive rate). The threshold is dependent on sample size and the number of segregating sites. This step is not usually undertaken in the case of D, which is frequently taken as supporting rejection of a null hypothesis of neutrality when its value has the property |D| *>* 2. However, our empirical analysis of the *ACKR1* gene region suggests that choosing this criterion for D rather than more precise thresholds significantly reduces its power (cf. Simonsen et al., 1995). We therefore obtained more accurate limits for D by simulation, as was done for *ρ*. Confidence limits for D can be obtained by conditioning on *θ* (as in Simonsen et al., 1995) or on *S*_*n*_ (as in Braverman et al., 1995). We chose the latter, since the model used to define *ρ*, and hence the thresholds used for *ρ*, were conditioned on *S*_*n*_. These thresholds are less conservative, and the resulting tests using D therefore more powerful, than those conditioned on *θ* as can be seen by comparing confidence limits shown in Supplementary Information Section 5 with those in Simonsen et al. (1995).

Our method can be used to test for departures from a demographic history characterised by constant population size, where we assume the absence of selection. It is effective in cases such as simple exponential growth (Figure 4), but can fail to detect departure from neutrality in some cases of more complex demographic history involving population bottlenecks (results not shown).

Unlike D, *ρ* depends on the identification of the ancestral states of the single nucleotide polymorphisms (SNPs) at segregating sites. When using *ρ*, consideration therefore needs to be given to the degree of confidence once has in the ascription of ancestral alleles to SNPs. Potential mitigations include confidence testing using alternate ancestral alleles; corrections to known biases in the use of parsimony to infer ancestral alleles (Hernandez et al., 2007; Keightley and Jackson, 2018); and deleting ambiguous SNPs from the data set. Because *ρ* is quite robust to a lower number of segregating sites, the cost of data lost by adopting the latter course may be acceptable.

The gene *ACKR1* provided an example of a locus where there is a widely accepted biological case that selection has occurred, yet previous genome scans using D failed to support this. Our results using *ρ* strongly suggest non-neutrality in both African and east Asian populations. The values of D shown at Table 2 also support this in most cases, when used with specifically calculated thresholds. This exemplifies how useful information can be lost when the crude |D| *>* 2 threshold is used. When we compared the likelihood of the selection scenario proposed by Hamblin et al. (2002) at this locus directly to the likelihood of a neutral model, the Hamblin scenario was preferred in the African populations by odds ratios *>* 5. The FY*O (rs2814778 T/C) variant is the only one in this genomic region with significant differences in allele frequencies between the African and east Asian populations. However, when the gene region was divided into four sub-regions, the signal for selection in African populations was weakest in the sub-region containing the FY*O variant. This result may be anomalous, nevertheless there is no support for this being a region of relatively high selection within the gene locus. On the other hand, both *ρ* and D identify this as the sub-region with the strongest evidence for selection in the Asian population (cf. Hamblin et al., 2002).

The most widely supported hypothesis to explain the divergence of African populations at this locus is that positive selection for the FY*O variant has occurred in African populations. This is supported by the determination of the T allele, which is almost universally present in non-African populations, as ancestral, based on chimpanzee (*Pan troglodytes*) and macaque (*Macaca mulatta*) outgroups. It could be argued that this phylogenetic inference is compromised by a lack of independence: chimpanzees, macaques and other primates are potentially susceptible to vivax malaria or its near relatives (Kaiser et al., 2010; Escalante et al., 2005). Indeed, it has been proposed that human *P. vivax* is a result of inter-species transmission from macaques in east Asia (Escalante et al., 2005; Mu et al., 2005). An alternative hypothesis to that of positive selection for FY*O in human populations in Africa would be that the C allele is ancestral and that selection favouring the T allele occurred in human as well as primate populations in east Asia. However, values of *ρ* calculated for chimpanzee and macaque populations are -1.78 and -0.26 respectively (see Materials and Methods for data sources). These values are considerably less than those we found in the human populations from Africa and so do not provide evidence against the hypothesis of positive selection for the FY*O variant in human populations from Africa.

Because *ρ* approximates the log-ratio of the average likelihood of all evolutionary models to that of the null model, it can be used directly to compare the degree of evidence for selection between data sets from different populations. We used this approach in our heat map analysis of chromosome band 2q11.1, with the aim of localising signals of selection in this region. Such a method is necessarily interrogative, rather than decisive, as comparisons between populations are affected by differences in sampling error. In many cases, the relative signals between loci obtained using *ρ* are similar to those obtained using D, as can be seen by comparing Supplementary Figure 5 to a similar analysis using D shown in Figure S2. However, for the region chr2:96845244-96865244 (A in Figure 5), *ρ* reveals a signal of selection that is less detectable using D. This is consistent with our simulation results demonstrating greater power for *ρ*. Overall, our analysis assisted in the identification of gene and regulatory feature loci in this region showing strong signals of selection.

In making inferences from the SFS, the impact of errors that may have been introduced into the data by upstream processes such as sequence read mapping and sequencing should be considered, particularly as sample sizes increase (Keinan and Clark, 2012). Errors are introduced in these processes to varying degrees and may bias results in different directions, depending on the bioinformatic methods used, e.g. direct sequencing or direct estimation of SFS. In any investigation, initial decisions need to be made on trade-offs between cost and choices of sample size, the length of genome to sequence, sequencing coverage and the choice of a bioinformatic pipeline. Direct maximum likelihood estimation of the SFS from aligned sequencing reads has been recommended for neutrality testing on the basis that it introduces less bias into estimation of the SFS and derived statistics than allele frequency estimation based on individual genotype calls (Han et al., 2013). A further issue with multi-sample genotype calling in particular is that software packages such as GATK and SAMtools use a prior based on the Wright-Fisher distribution, which will bias tests of neutrality (Han et al., 2013). Direct estimation of the SFS is implemented in software such as ANGSD (Korneliussen et al., 2014), which estimates the SFS using an expectation–maximization method (Li, 2011; Nielsen et al., 2012). However, even if an unbiased estimation of the SFS is used, the estimate will still have variability and ignoring this may potentially bias evaluation of the null hypothesis. This could be overcome if a posterior distribution of the SFS could be obtained, which could be integrated over equation (8), but the authors are not aware of any tools that generate such a distribution.

The major impact of sequencing error on the SFS, particularly for low-coverage (*<* 10 ×) data, is at sites with low allele frequency, such as singletons and doubletons. Such error will bias subsequent tests such as D (Han et al., 2013). For this reason, it has been recommended that low-frequency variants be removed from the analysis altogether (Achaz, 2009; Ferretti et al., 2013). In the case of D, this has been shown to reduce bias due to sequencing error (Han et al., 2013). Our method can adapt to the removal of low frequency variants by a modification of the likelihood equation (8). Details are given in Supplementary Information Section 7.

In summary, we have described a test for departure from selective neutrality that derives its power from the likelihood principle. While being relatively computationally intensive, it is powerful compared to alternatives and flexible in its ability to adapt to different null hypotheses or to account for potential sequencing error.

## ACKNOWLEDGMENTS

This research was supported by an Australian Government Research Training Program (RTP) Scholarship to HS. The authors would like to thank John Wakeley for helpful comments.

## REFERENCES

1000 Genomes Project Consortium; Auton, A., Brooks, L., Durbin, R., Garrison, E., Kang, H., Korbel, J., Marchini, J., McCarthy, S., McVean, G., and Abecasis, G. (2015). A global reference for human genetic variation. Nature, 526(7571):68–74. [ftp://ftp.1000genomes.ebi.ac.uk/vol1/ftp/release/20130502/].

Achaz, G. (2009). Frequency spectrum neutrality tests: one for all and all for one. Genetics, 183(1):249–258.

Akey, J. M., Eberle, M. A., Rieder, M. J., Carlson, C. S., Shriver, M. D., Nickerson, D. A., and Kruglyak, L. (2004). Population history and natural selection shape patterns of genetic variation in 132 genes. PLoS Biology, 2(10):e286.

Altshuler, D. and Donnelly, P. (2005). A haplotype map of the human genome. Nature, 437(7063):1299–1320.

Bhaskar, A. and Song, Y. S. (2014). Descartes’ rule of signs and the identifiability of population demographic models from genomic variation data. Annals of Statistics, 42(6):2469–2493.

Bimber, B. N., Yan, M. Y., Peterson, S. M., and Ferguson, B. (2019). mGAP: the macaque genotype and phenotype resource, a framework for accessing and interpreting macaque variant data, and identifying new models of human disease. BMC Genomics, 20(1):1–9. Funding from NIH R24OD021324. https://mgap.ohsu.edu.

Braverman, J. M., Hudson, R. R., Kaplan, N. L., Langley, C. H., and Stephan, W. (1995). The hitchhiking effect on the site frequency spectrum of DNA polymorphisms. Genetics, 140(2):783–796.

Carter, R. (2003). Speculations on the origins of Plasmodium vivax malaria. Trends in Parasitology, 19(5):214–219.

Casbon, J. et al. (2012). PyVCF. [https://pypi.org/project/PyVCF/].

Charlesworth, B., Morgan, M., and Charlesworth, D. (1993). The effect of deleterious mutations on neutral molecular variation. Genetics, 134(4):1289–1303.

Corbett, M. A., Kroes, T., Veneziano, L., Bennett, M. F., Florian, R., Schneider, A. L., Coppola, A., Licchetta, L., Franceschetti, S., Suppa, A., et al. (2019). Intronic ATTTC repeat expansions in STARD7 in familial adult myoclonic epilepsy linked to chromosome 2. Nature Communications, 10(1):1–10.

Cvijović, I., Good, B. H., and Desai, M. M. (2018). The effect of strong purifying selection on genetic diversity. Genetics, 209(4):1235–1278.

DeTemple, D. and Webb, W. (2014). Combinatorial reasoning: An introduction to the art of counting. John Wiley & Sons, Hoboken, New Jersey.

Escalante, A. A., Cornejo, O. E., Freeland, D. E., Poe, A. C., Durrego, E., Collins, W. E., and Lal, A. A. (2005). A monkey’s tale: the origin of Plasmodium vivax as a human malaria parasite. Proceedings of the National Academy of Sciences, 102(6):1980–1985.

Ewing, G. and Hermisson, J. (2010). MSMS: a coalescent simulation program including recombination, demographic structure and selection at a single locus. Bioinformatics, 26(16):2064–2065.

Fay, J. C. and Wu, C.-I. (2000). Hitchhiking under positive Darwinian selection. Genetics, 155(3):1405–1413.

Ferretti, L., Ramos-Onsins, S. E., and Pérez-Enciso, M. (2013). Population genomics from pool sequencing. Molecular Ecology, 22(22):5561–5576.

Fu, Y.-X. (1995). Statistical properties of segregating sites. Theoretical Population Biology, 48(2):172–197.

Fu, Y.-X. and Li, W.-H. (1993). Statistical tests of neutrality of mutations. Genetics, 133(3):693–709.

Grandi, N., Cadeddu, M., Blomberg, J., and Tramontano, E. (2016). Contribution of type W human endogenous retroviruses to the human genome: characterization of HERV-W proviral insertions and processed pseudogenes. Retrovirology, 13(1):1–25.

Griffiths, R. and Tavaré, S. (1994). Sampling theory for neutral alleles in a varying environment. Philosophical Transactions of the Royal Society of London B: Biological Sciences, 344(1310):403–410.

Griffiths, R. C. and Tavaré, S. (1998). The age of a mutation in a general coalescent tree. Stochastic Models, 14(1-2):273–295.

Hamblin, M. T. and Di Rienzo, A. (2000). Detection of the signature of natural selection in humans: evidence from the Duffy blood group locus. The American Journal of Human Genetics, 66(5):1669–1679.

Hamblin, M. T., Thompson, E. E., and Rienzo, A. D. (2002). Complex signatures of natural selection at the Duffy blood group locus. The American Journal of Human Genetics, 70(2):369–383.

Han, E., Sinsheimer, J. S., and Novembre, J. (2013). Characterizing bias in population genetic inferences from low-coverage sequencing data. Molecular Biology and Evolution, 31(3):723–735.

Hein, J., Schierup, M., and Wiuf, C. (2005). Gene genealogies, variation and evolution: a primer in coalescent theory. Oxford University Press, Oxford UK.

Hernandez, R. D., Williamson, S. H., and Bustamante, C. D. (2007). Context dependence, ancestral misidentification, and spurious signatures of natural selection. Molecular Biology and Evolution, 24(8):1792–1800.

Hodgson, J. A., Pickrell, J. K., Pearson, L. N., Quillen, E. E., Prista, A., Rocha, J., Soodyall, H., Shriver, M. D., and Perry, G. H. (2014). Natural selection for the Duffy-null allele in the recently admixed people of Madagascar. Proceedings of the Royal Society B: Biological Sciences, 281(1789):20140930.

Hudson, R. R. (2002). Generating samples under a Wright-Fisher neutral model of genetic variation. Bioinformatics, 18(2):337–338.

Hunter, J. D. (2007). Matplotlib: A 2D graphics environment. Computing in Science & Engineering, 9(3):90–95.

Huttley, G. (2016). scitrack 0.1.1. [https://pypi.org/project/scitrack/0.1.1/].

Innan, H. and Stephan, W. (2003). Distinguishing the hitchhiking and background selection models. Genetics, 165(4):2307–2312.

Jeffreys, H. (1998). The theory of probability. Oxford University Press, Oxford, United Kingdom.

Josephs, E. B. and Wright, S. I. (2016). On the trail of linked selection. PLoS Genetics, 12(8):e1006240.

Kaiser, M., Löwa, A., Ulrich, M., Ellerbrok, H., Goffe, A. S., Blasse, A., Zommers, Z., Couacy-Hymann, E., Babweteera, F., Zuberbühler, K., et al. (2010). Wild chimpanzees infected with 5 Plasmodium species. Emerging Infectious Diseases, 16(12):1956– 1959.

Keightley, P. D. and Jackson, B. C. (2018). Inferring the probability of the derived vs. the ancestral allelic state at a polymorphic site. Genetics, 209(3):897–906.

Keinan, A. and Clark, A. G. (2012). Recent explosive human population growth has resulted in an excess of rare genetic variants. Science, 336(6082):740–743.

Kelleher, J., Etheridge, A. M., and McVean, G. (2016). Efficient coalescent simulation and genealogical analysis for large sample sizes. PLoS Computational Biology, 12(5):e1004842.

Kelleher, J., Thornton, K. R., Ashander, J., and Ralph, P. L. (2018). Efficient pedigree recording for fast population genetics simulation. PLoS Computational Biology, 14(11):e1006581.

Kelley, J. L., Madeoy, J., Calhoun, J. C., Swanson, W., and Akey, J. M. (2006). Genomic signatures of positive selection in humans and the limits of outlier approaches. Genome Research, 16(8):980–989.

Kimura, M. (1969). The number of heterozygous nucleotide sites maintained in a finite population due to steady flux of mutations. Genetics, 61(4):893–903.

Kimura, M. (1983). The neutral theory of molecular evolution. Cambridge University Press, Cambridge, UK.

Kingman, J. F. C. (1982a). The coalescent. Stochastic Processes and their Applications, 13(3):235–248.

Kingman, J. F. C. (1982b). On the genealogy of large populations. Journal of Applied Probability, 19(A):27–43.

Knight, R., Maxwell, P., Birmingham, A., Carnes, J., Caporaso, J. G., Easton, B. C., Eaton, M., Hamady, M., Lindsay, H., Liu, Z., et al. (2007). Pycogent: a toolkit for making sense from sequence. Genome Biology, 8(8):1–16.

Korneliussen, T. S., Albrechtsen, A., and Nielsen, R. (2014). ANGSD: analysis of next generation sequencing data. BMC Bioinformatics, 15(1):1–13.

Lenders, J. W., Eisenhofer, G., Mannelli, M., and Pacak, K. (2005). Phaeochromocytoma. The Lancet, 366(9486):665–675.

Li, H. (2011). A statistical framework for SNP calling, mutation discovery, association mapping and population genetical parameter estimation from sequencing data. Bioinformatics, 27(21):2987–2993.

Li, H., Handsaker, B., Wysoker, A., Fennell, T., Ruan, J., Homer, N., Marth, G., Abecasis, G., and Durbin, R. (2009). The sequence alignment/map format and SAMtools. Bioinformatics, 25(16):2078–2079.

Livingstone, F. B. (1984). The Duffy blood groups, vivax malaria, and malaria selection in human populations: a review. Human Biology, 56(3):413–425.

McKinney, W. (2010). Data structures for statistical computing in Python. In van der Walt, S.and Millman, J., editors, Proceedings of the 9th Python in Science Conference, pages 51–56.

Miller, L. H., Mason, S. J., Clyde, D. F., and McGinniss, M. H. (1976). The resistance factor to Plasmodium vivax in blacks: the Duffy-blood-group genotype, FyFy. New England Journal of Medicine, 295(6):302–304.

Mooers, A. O. and Heard, S. B. (1997). Inferring evolutionary process from phylogenetic tree shape. The Quarterly Review of Biology, 72(1):31–54.

Mu, J., Joy, D. A., Duan, J., Huang, Y., Carlton, J., Walker, J., Barnwell, J., Beerli, P., Charleston, M. A., Pybus, O. G., and Su, X.-z. (2005). Host switch leads to emergence of Plasmodium vivax malaria in humans. Molecular Biology and Evolution, 22(8):1686–1693.

Nielsen, R., Hubisz, M. J., Hellmann, I., Torgerson, D., Andrés, A. M., Albrechtsen, A., Gutenkunst, R., Adams, M. D., Cargill, M., Boyko, A., Indap, A., Bustamante, C. D., and Clark, A. G. (2009). Darwinian and demographic forces affecting human protein coding genes. Genome Research, 19(5):838–849.

Nielsen, R., Korneliussen, T., Albrechtsen, A., Li, Y., and Wang, J. (2012). SNP calling, genotype calling, and sample allele frequency estimation from new-generation sequencing data. PLoS ONE, 7(7):e37558.

Prado-Martinez, J., Sudmant, P. H., Kidd, J. M., Li, H., Kelley, J. L., Lorente-Galdos, B., Veeramah, K. R., Woerner, A. E., O’Connor, T. D., Santpere, G., et al. (2013). Great ape genetic diversity and population history. Nature, 499(7459):471–475. http://biologiaevolutiva.org/greatape.

Qin, Y., Yao, L., King, E. E., Buddavarapu, K., Lenci, R. E., Chocron, E. S., Lechleiter, J. D., Sass, M., Aronin, N., Schiavi, F., Boaretto, F., Opocher, G., Toledo, R. A., Toledo, S. P. A., Stiles, C., Aguiar, R. C. T., and Dahia, P. L. M. (2010). Germline mutations in TMEM127 confer susceptibility to pheochromocytoma. Nature Genetics, 42(3):229–33.

Robert, C. P., Casella, G., and Casella, G. (2010). Introducing Monte Carlo methods with R, volume 18. Springer, New York.

Ronacher, A. (2009). click 7.0. [https://pypi.org/project/click/].

Rudin, W. (1987). Real and complex analysis. McGraw-Hill, New York.

Sabeti, P. C., Schaffner, S. F., Fry, B., Lohmueller, J., Varilly, P., Shamovsky, O., Palma, A., Mikkelsen, T., Altshuler, D., and Lander, E. (2006). Positive natural selection in the human lineage. Science, 312(5780):1614–1620.

Sainudiin, R., Thornton, K., Harlow, J., Booth, J., Stillman, M., Yoshida, R., Griffiths, R., Gil, M., and Donnelly, P. (2011). Experiments with the site frequency spectrum. Bulletin of Mathematical Biology, 73(4):829–872.

Simonsen, K. L., Churchill, G. A., and Aquadro, C. F. (1995). Properties of statistical tests of neutrality for DNA polymorphism data. Genetics, 141(1):413–429.

Slatkin, M. and Hudson, R. R. (1991). Pairwise comparisons of mitochondrial-DNA sequences in stable and exponentially growing populations. Genetics, 129(2):555– 562.

Sloane, N. J. et al. (2003). The on-line encyclopedia of integer sequences, published electronically at https://oeis.org, accessed 13/07/2016.

Stajich, J. E. and Hahn, M. W. (2004). Disentangling the effects of demography and selection in human history. Molecular Biology and Evolution, 22(1):63–73.

Tajima, F. (1989). Statistical method for testing the neutral mutation hypothesis by DNA polymorphism. Genetics, 123(3):585–595.

Tretyakov, K. (2013). pyliftover 0.4. [https://pypi.org/project/pyliftover/].

Virtanen, P., Gommers, R., Oliphant, T. E., Haberland, M., Reddy, T., Cournapeau, D., Burovski, E., Peterson, P., Weckesser, W., Bright, J., et al. (2020). Scipy 1.0: fundamental algorithms for scientific computing in python. Nature Methods, 17(3):261–272.

Vitti, J. J., Grossman, S. R., and Sabeti, P. C. (2013). Detecting natural selection in genomic data. Annual Review of Genetics, 47:97–120.

Waskom, M., Botvinnik, O., O’Kane, D., Hobson, P., Lukauskas, S., Gemperline, D. C., Augspurger, T., Halchenko, Y., Cole, J. B., Warmenhoven, J., de Ruiter, J., Pye, C., Hoyer, S., Vanderplas, J., Villalba, S., Kunter, G., Quintero, E., Bachant, P., Martin, M., Meyer, K., Miles, A., Ram, Y., Yarkoni, T., Williams, M. L., Evans, C., Fitzgerald, C., Brian Fonnesbeck, C., Lee, A., and Qalieh, A. (2017). Seaborn: v0.8.1. https://doi.org/10.5281/zenodo.883859.

Zhao, Y., Cao, X., Guo, M., Wang, X., Yu, T., Ye, L., Han, L., Hei, L., Tao, W., Tong, Y., Xu, Y., and Zhong, J. (2018). NEURL3 is an inducible antiviral effector to inhibit HCV assembly by targeting viral E1 glycoprotein. Journal of Virology, 92(21):e01123–18.

