## Supplementary Information for "A New Likelihood-based Test for Natural Selection"

#### 1 An example: tree topology is not equivalent to TPC

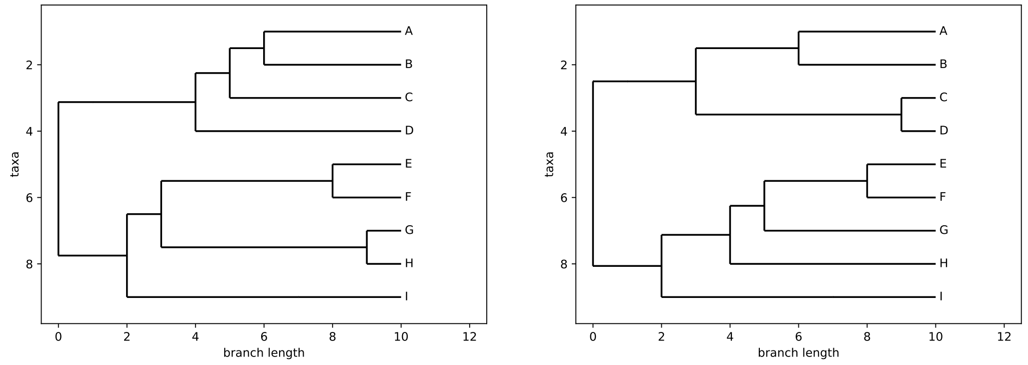


(a) Two different tree topologies

$$\begin{bmatrix} 0 & 0 & 0 & 1 & 1 & 0 & 0 & 0 \\ 1 & 0 & 0 & 2 & 0 & 0 & 0 & 0 \\ 1 & 2 & 0 & 1 & 0 & 0 & 0 & 0 \\ 2 & 2 & 1 & 0 & 0 & 0 & 0 & 0 \\ 3 & 3 & 0 & 0 & 0 & 0 & 0 & 0 \\ 5 & 2 & 0 & 0 & 0 & 0 & 0 & 0 \\ 7 & 1 & 0 & 0 & 0 & 0 & 0 & 0 \\ 9 & 0 & 0 & 0 & 0 & 0 & 0 & 0 \end{bmatrix}$$

(b) Common tree matrix

Figure S1: Subfigure (a) shows two trees that have the same TPC or tree matrix, but different rooted topologies. (b) The common tree matrix for the two trees. The levels of the tree are represented by the rows of the matrix: reading from top to bottom corresponds to tracing from the root to the leaves.

#### 2 Inverse of the matrix $A_n$

We have denoted by $A_n$ the $n-1 \times n-1$ matrix corresponding to equation (4) in Materials and Methods as well as to equation (3.3) of Griffiths and Tavaré (1998).

From equations (4) and (3) we have:

$$A_n[i,j] = (i+1)\frac{\binom{n-j-1}{i-1}}{\binom{n-1}{i}} \quad \text{if } j \leq n-i\text{; 0 otherwise}$$

We show that the inverse of $A_n$ is given by $B_n$ where:

$$B_n[i,j] = (-1)^{i+j-n}\binom{n-1}{j}\binom{j-1}{n-i-1}/(j+1) \quad \text{if } j \geq n-i\text{; 0 otherwise.}$$

Let $C_n = A_n B_n$. Then

$$C_n[i,j] = \sum_{k=1}^{n-1} A_n[i,k]B_n[k,j]$$

All terms are zero if $k > n-i$ or $k < n-j$. If $i > j$, one of these alternatives always holds, so for $i > j$, $C_n[i,j] = 0$.

If $i = j$, all terms are zero except for $k = n-i$, hence:

$$C_n[i,i] = \frac{(i+1)\binom{i-1}{i-1}}{\binom{n-1}{i}}(-1)^0\binom{n-1}{i}\frac{\binom{i-1}{i-1}}{i+1} = 1$$

If $j > i$ we have, excluding zero terms:

$$
\begin{aligned}
C_n[i,j] &= \sum_{k=n-j}^{n-i} (i+1)\frac{\binom{n-k-1}{i-1}}{\binom{n-1}{i}}(-1)^{k+j-n}\binom{n-1}{j}\binom{j-1}{n-k-1}/(j+1) \\
&= \left(\frac{i+1}{j+1}\right)\frac{\binom{n-1}{j}}{\binom{n-1}{i}}\sum_{k=n-j}^{n-i}(-1)^{k+j-n}\binom{n-k-1}{i-1}\binom{j-1}{n-k-1} \\
&= \left(\frac{i+1}{j+1}\right)\frac{\binom{n-1}{j}}{\binom{n-1}{i}}\sum_{k=n-j}^{n-i}(-1)^{k+j-n}\frac{(n-k-1)!}{(i-1)!(n-k-i)!}\frac{(j-1)!}{(n-k-1)!(j-n+k)!} \\
&= \left(\frac{i+1}{j+1}\right)\frac{\binom{n-1}{j}}{\binom{n-1}{i}}\frac{(j-1)!}{(i-1)!}\sum_{k=n-j}^{n-i}(-1)^{k+j-n}\frac{1}{(n-k-i)!(j-n+k)!} \\
&= \left(\frac{i+1}{j+1}\right)\frac{\binom{n-1}{j}}{\binom{n-1}{i}}\frac{(j-1)!}{(i-1)!}\frac{1}{(j-i)!}\sum_{k=n-j}^{n-i}(-1)^{k+j-n}\binom{j-i}{k-n+j} \\
&= \frac{i+1}{j+1}\frac{\binom{n-1}{j}}{\binom{n-1}{i}}\frac{(j-1)!}{(i-1)!}\frac{(j-1)!}{(i-1)!}\frac{1}{(j-i)!}\sum_{m=0}^{j-i}(-1)^m\binom{j-i}{m} \quad \text{(setting } m = k-n+j\text{)} \\
&= \frac{i+1}{j+1}\frac{\binom{n-1}{j}}{\binom{n-1}{i}}\frac{(j-1)!}{(i-1)!}\frac{1}{(j-i)!}(1-1)^{j-i} \\
&= 0
\end{aligned}
$$

### 3 1000 Genomes Project populations

The following are the 26 populations in the 1KG Project dataset used in this paper, with population and super-population codes.

```
CHB/EAS Han Chinese in Beijing, China
JPT/EAS Japanese in Tokyo, Japan
CHS/EAS Southern Han Chinese
CDX/EAS Chinese Dai in Xishuangbanna, China
KHV/EAS Kinh in Ho Chi Minh City, Vietnam

CEU/EUR Utah Residents with Northern and Western European Ancestry
TSI/EUR Toscani in Italia
FIN/EUR Finnish in Finland
GBR/EUR British in England and Scotland
IBS/EUR Iberian Population in Spain

YRI/AFR Yoruba in Ibadan, Nigeria
LWK/AFR Luhya in Webuye, Kenya AFR
GWD/AFR Gambian in Western Divisions in the Gambia
MSL/AFR Mende in Sierra Leone
ESN/AFR Esan in Nigeria
ASW/AFR Americans of African Ancestry in SW USA
ACB/AFR African Caribbeans in Barbados

MXL/AMR Mexican Ancestry from Los Angeles USA
PUR/AMR Puerto Ricans from Puerto Rico
CLM/AMR Colombians from Medellin, Colombia
PEL/AMR Peruvians from Lima, Peru

GIH/SAS Gujarati Indian from Houston, Texas
PJL/SAS Punjabi from Lahore, Pakistan
BEB/SAS Bengali from Bangladesh
STU/SAS Sri Lankan Tamil from the UK
ITU/SAS Indian Telugu from the UK
```

### 4 Bottleneck scenario for European populations

This section contains details of the bottleneck demographic scenario used as the null scenario for European populations in the analysis of a region from Human chromosome band 2q11.1 (refer Figure 5). The scenario is as proposed in (Stajich and Hahn, 2004). Output is as generated by the msprime package (Kelleher et al., 2016). Note that "start" and "end" refer to the effective population size at the start and end of each epoch. Epochs are defined in generations counting back in time from the present.

```
==============================
Epoch: 0 -- 500.0 generations
==============================
     start     end      growth_rate |     0
   -------- --------       -------- | --------
0 | 6.6e+03  6.6e+03              0 |     0

Events @ generation 500.0
   - Population parameter change for -1: initial_size -> 3300.0 growth_rate -> 0
===================================
Epoch: 500.0 -- 1500.0 generations
===================================
     start     end      growth_rate |     0
   -------- --------       -------- | --------
0 | 3.3e+03  3.3e+03              0 |     0

Events @ generation 1500.0
   - Population parameter change for -1: initial_size -> 10000.0 growth_rate -> 0
=================================
Epoch: 1500.0 -- inf generations
=================================
     start     end      growth_rate |     0
   -------- --------       -------- | --------
0 |  1e+04    1e+04               0 |     0
```

### 5 Thresholds calculated for Tajima's D

| n = | 10 | 20 | 50 | 75 | 100 |
|---|---|---|---|---|---|
| **Lower bound** | -1.73 | -1.73 | -1.67 | -1.65 | -1.63 |
| **Upper bound** | 1.73 | 1.81 | 1.92 | 1.95 | 1.99 |

(a) $\alpha = 0.05$

| | | | | | |
|---|---|---|---|---|---|
| **Lower bound** | -1.53 | -1.52 | -1.48 | -1.47 | -1.45 |
| **Upper bound** | 1.45 | 1.51 | 1.56 | 1.61 | 1.62 |

(b) $\alpha = 0.1$

Table S1: Thresholds for Tajima's D used for Figure 2 (positive selection) for a range of values of $n$. Subtables record different significance levels $\alpha$. The level $\alpha = 0.1$ is is used for the one-sided test. For all cases $S_n = 10$.

| $\mathbf{S_n}$ = | 65 | 100 | 125 | 150 | 200 | 250 | 325 | 400 |
|---|---|---|---|---|---|---|---|---|
| **Lower bound** | -1.73 | -1.73 | -1.72 | -1.74 | -1.73 | -1.72 | -1.72 | -1.73 |
| **Upper bound** | 1.71 | 1.69 | 1.68 | 1.66 | 1.67 | 1.67 | 1.67 | 1.67 |

(a) $\alpha = 0.05$

| | | | | | | | | |
|---|---|---|---|---|---|---|---|---|
| **Lower bound** | -1.48 | -1.49 | -1.49 | -1.48 | -1.48 | -1.49 | -1.49 | -1.48 |
| **Upper bound** | 1.41 | 1.41 | 1.40 | 1.41 | 1.39 | 1.39 | 1.39 | 1.39 |

(b) $\alpha = 0.1$

Table S2: Thresholds for Tajima's D used for Figure 3 (negative selection) for a range of values of $S_n$. Subtables record different significance levels $\alpha$. The level $\alpha = 0.1$ is is used for the one-sided test. In all cases $n = 20$.

### 6 Analysis of 2q11.1 region using Tajima's D

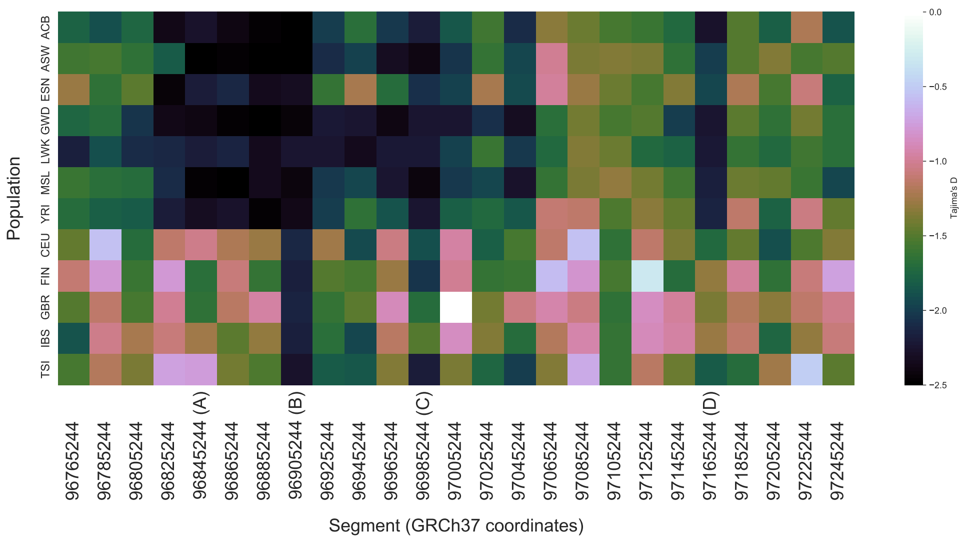


Figure S2: Values of Tajima's D by 20-kb segment and population for the 2q11.1 region.

### 7 Reducing bias due to sequencing error

We show how to take account of the potential for sequencing error by modifying the likelihood equation (8) in Materials and Methods. If we introduce the potential for error in the SFS due to sequencing errors, the true number of segregating sites will have an unknown value, which we denote by $S_n^{\mathcal{T}}$. We will remove from the analysis variants at frequencies $1 \leq i \leq m$, for some $m < n-1$. The likelihood given the truncated SFS $s_{m+1}, \ldots, s_{n-1}$ is given by:

$$\mathcal{L}(\mathbf{q}|\mathbf{s}') = \mathbb{P}(\mathbf{s}'|\mathbf{q}) = p_{mult}(\mathbf{s}', S_n^{\mathcal{T}}, \mathbf{q}') \tag{1}$$

where:

$$\mathbf{q}' = \left(1 - \sum_{i=m+1}^{n-1} q_i, q_{m+1}, \ldots, q_{n-1}\right) \text{ ; and}$$
$$\mathbf{s}' = \left(S_n^{\mathcal{T}} - \sum_{i=m+1}^{n-1} s_i, s_{m+1}, \ldots, s_{n-1}\right)$$

That is, the new likelihood is a multinomial distribution with parameters $S_n^{\mathcal{T}}$ as the number of draws, $\mathbf{q}'$ as the vector of $n-m$ probabilities and $\mathbf{s}'$ as the outcome. The likelihood is integrated over the distribution of the probability vectors $\mathbf{q}$ as before. We can estimate $S_n^{\mathcal{T}}$ by (Ferretti et al., 2013):

$$\widehat{S_n^{\mathcal{T}}} = \sum_s p(s)$$

where summation is over all sites $s$ and $p(s)$ is the probability that a SNP call at the site $s$ is true, derived from the quality score determined by the SNP calling software platform used or by third-party software post-sequencing. $\widehat{S_n^{\mathcal{T}}}$ would need to be rounded to the nearest integer. A modification to avoid the need for such rounding and better reflect the variability in $\widehat{S_n^{\mathcal{T}}}$ would be to draw samples of its sampling distribution, namely a Poisson binomial distribution for which the parameters are the vectors $p(s)$. The likelihood in equation (1) would then require Monte Carlo integration over this distribution on $S_n^{\mathcal{T}}$ as well as over the probability vectors $\mathbf{q}$.

### References

Ferretti, L., Ramos-Onsins, S. E., and Pérez-Enciso, M. (2013). Population genomics from pool sequencing. *Molecular Ecology*, 22(22):5561–5576.

Griffiths, R. C. and Tavaré, S. (1998). The age of a mutation in a general coalescent tree. *Stochastic Models*, 14(1-2):273–295.

Kelleher, J., Etheridge, A. M., and McVean, G. (2016). Efficient coalescent simulation and genealogical analysis for large sample sizes. *PLoS Computational Biology*, 12(5):1–22.

Stajich, J. E. and Hahn, M. W. (2004). Disentangling the effects of demography and selection in human history. *Molecular Biology and Evolution*, 22(1):63–73.
